# *Cancer Cachexia in STK11/LKB1*-mutated NSCLC is Dependent on Tumor-secreted GDF15

**DOI:** 10.1101/2024.06.14.598891

**Authors:** Jinhai Yu, Tong Guo, Arun Gupta, Ernesto M. Llano, Naureen Wajahat, Sean Slater, Qing Deng, Esra A. Akbay, John M. Shelton, Bret M. Evers, Zhidan Wu, Iphigenia Tzameli, Evanthia Pashos, John D. Minna, Puneeth Iyengar, Rodney E. Infante

**Affiliations:** Center for Human Nutrition, University of Texas Southwestern Medical Center, Dallas, Texas, USA; Harold C. Simmons Comprehensive Cancer Center, University of Texas Southwestern Medical Center, Dallas, Texas, USA; Department of Internal Medicine, University of Texas Southwestern Medical Center, Dallas, Texas, USA; Department of Pathology, University of Texas Southwestern Medical Center, Dallas, Texas, USA; Department of Pharmacology, University of Texas Southwestern Medical Center, Dallas, Texas, USA; Hamon Center for Therapeutic Oncology Research, University of Texas Southwestern Medical Center, Dallas, Texas, USA; Department of Radiation Oncology, University of Texas Southwestern Medical Center, Dallas, Texas, USA; Department of Molecular Genetics, University of Texas Southwestern Medical Center, Dallas, Texas, USA; Department of Radiation Oncology, Memorial Sloan Kettering Cancer Center, New York, New York, USA; Druckenmiller Center for Lung Cancer Research, Memorial Sloan Kettering Cancer Center, New York, New York, USA; Internal Medicine Research Unit, Pfizer Inc., Cambridge, MA, USA

**Keywords:** STK11/LKB1, GDF15, non-small cell lung cancer, cachexia, wasting, adipose loss, muscle loss, weight loss, anorexia

## Abstract

Cachexia is a wasting syndrome comprised of adipose, muscle, and weight loss observed in cancer patients. Tumor loss-of-function mutations in *STK11/LKB1*, a regulator of the energy sensor AMP-activated protein kinase, induce cancer cachexia (CC) in preclinical models and are associated with cancer-related weight loss in NSCLC patients. Here we characterized the relevance of the NSCLC-associated cachexia factor growth differentiation factor 15 (GDF15) in several patient-derived and genetically engineered *STK11/LKB1*-mutant NSCLC cachexia lines. Both tumor mRNA expression and serum concentrations of tumor-derived GDF15 were significantly elevated in multiple mice transplanted with patient-derived *STK11/LKB1*-mutated NSCLC lines. GDF15 neutralizing antibody administered to mice transplanted with patient- or mouse-derived *STK11/LKB1*-mutated NSCLC lines suppressed cachexia-associated adipose loss, muscle atrophy, and changes in body weight. The silencing of *GDF15* in multiple human NSCLC lines was also sufficient to eliminate *in vivo* circulating GDF15 levels and abrogate cachexia induction, suggesting that tumor and not host tissues represent a key source of GDF15 production in these cancer models. Finally, reconstitution of wild-type *STK11/LKB1* in a human *STK11/LKB1* loss-of-function NSCLC line that normally induces cachexia *in vivo* correlated with the absence of tumor-secreted GDF15 and rescue from the cachexia phenotype. The current data provide evidence for tumor-secreted GDF15 as a conduit and a therapeutic target through which NSCLCs with *STK11/LKB1* loss-of-function mutations promote cachexia-associated wasting.

## INTRODUCTION

Despite long-standing clinical evidence and knowledge associating the wasting syndrome of cachexia with the development of certain cancers, there has been little progress in identifying cancer cachexia (CC) biomarkers and therapeutic interventions. This crucial knowledge gap has resulted from several factors including: 1) heterogeneity in CC mechanisms in patient populations with distinct primary cancers; 2) the presence of CC mechanisms involving multiple tissues; 3) previous “one-size-fits-all” approaches to treat CC by targeting individual molecules and pathways that were are not universally relevant across different primary cancer and host tissues; 4) efforts to only combat muscle atrophy and not adipose loss or anorectic signals to the brain; and 5) the lack of preclinical isogenic models that provide the most stringent controls for quantitative assessments of CC drivers and their roles as therapeutic targets.

To overcome some of these barriers in better understanding the fundamental mechanisms of CC, we reported a screen of human NSCLCs assessing their capacity to induce cachexia-associated fat and muscle wasting in immunodeficient mice.^1^ We dichotomized a cohort of patient-derived NSCLC lines with CC-inducing capacity *in vivo* and a matched cohort of NSCLC lines without CC-inducing capacity. Interestingly, 80% of the CC-inducing NSCLCs possessed loss-of-function variants in *STK11/LKB1*, the upstream regulator of the cellular energy sensor AMP-activated protein kinase (AMPK). *STK11/LKB1* was wild type in every line that failed to induce CC, suggesting that variants in the gene are a potential biomarker for CC. Interestingly, *STK11/LKB1* silencing in genetically engineered *Kras* mutant mice also caused the majority of animals to develop CC.^2^ We next showed that silencing of *STK11/LKB1* in a non-cachexia inducing NSCLC line converted it into one with CC-inducing potential *in vivo*, establishing that the functional absence of this molecule was directly responsible for the development of cachexia.^1^ Finally, in a cohort of nearly 250 advanced NSCLC patients, ctDNA-based liquid biopsy assays established a significant correlation between a patient having a tumor with *STK11/LKB1* functional variant and the presence of CC-associated wasting at cancer diagnosis.^1^

Having shown that tumor STK11/LKB1 loss-of-function is not only a predictive and prognostic biomarker but also functionally involved in CC, we sought to identify the downstream tumor-intrinsic molecules transducing mutant *STK11/LKB1* CC signals. A candidate approach evaluating tumor-secreted factors with the potential to influence host wasting that had a dependence on tumor STK11/LKB1 functional loss led us to investigate GDF15 as such a molecule. GDF15 is an inflammatory cytokine of the TGFβ super family with many pathophysiologic roles, including in metabolism where it is thought to promote anorexia and subsequent weight loss by direct signaling of the hindbrain through the glial-cell-derived neurotrophic factor family receptor α-like (GFRAL).^3–8^ Originally discovered to be released with tissue injury, GDF15 has been found to regulate a number of fundamental processes.^9, 10^ This secreted molecule first is produced as a pro-form of protein within cells and only after processing by proteases is GDF15 converted into a mature form that can be found in systemic circulation acting centrally on GFRAL-positive neurons in the hindbrain.^11, 12^ Systemically, GDF15-induced anorexia was previously only thought to be relevant when triggered by platinum-based and other cytotoxic chemotherapies offered to cancer patients.^13, 14^ The presumption supported by data has been that the elevated circulating concentrations of GDF15 are a consequence of normal tissue toxicities secondary to patient exposure to these cytotoxic agents or as a result of disease progression. Recent investigations also highlight a link between GDF15 induction and the development of NSCLC-associated and other cancer-associated cachexia.^15–17^ Independent of central effects, there are now other suggestions that GFD15 may signal peripherally during times of exercise in regulating contractility and skeletal muscle action.^18^ In addition, antibody/small molecule inhibitors of GDF15/GFRAL signaling are being assessed in clinical trials to block cancer-associated cachexia.^13, 19, 20^

Our fundamental understanding of whether circulating GDF15 primarily originates from tumor or host cells, and the dependencies of tumor systems on GDF15 action remain unknown. Here, we took advantage of our previous efforts to characterize the cachexia-inducing potential of a cohort of patient-derived NSCLC lines to delineate a tumor’s GDF15 dependence in inducing cachexia and in identifying tumor intrinsic (human) versus extrinsic (murine host) contributions to systemic GDF15 concentrations. We determined that out of 8 human *STK11/LKB1*-mutated, cachexia-inducing and 7 wild-type *STK11/LKB1,* non-cachexia-inducing NSCLC lines injected into immunodeficient mice to form tumors, there were significant differences in human (tumor) but not murine (host) *GDF15* mRNA levels and human (tumor) serum GDF15 concentrations between cohorts. Specifically, mice injected with the 8 human NSCLC cachexia-inducing lines displayed significantly elevated human serum GDF15 concentrations compared to the mice injected with the 7 human NSCLC non-cachexia inducing counterparts. Independently, both a neutralizing antibody against GDF15 and silencing of *GDF15* in tumor cells suppressed *STK11/LKB1*-mutated, NSCLC-associated cachexia, suggesting a GDF15-dependence. We also demonstrated that silencing of *STK11/LKB1* in a non-cachexia inducing patient-derived line led to the promotion of cachexia *in vivo*, which was subsequently suppressed in the setting of GDF15 antibody administration. In parallel, we verified that a *Kras*, *Trp53*, and *Stk11/Lkb1* mutant line (KPL) derived from a genetically-engineered mouse model also induces the host cachexia phenotype when injected into syngeneic mice, the latter similarly ameliorated with the administration of GDF15 antibody. Finally, reconstituting wild-type *STK11/LKB1* in a patient-derived NSCLC line with endogenously-mutated *STK11/LKB1* decreased tumor-secreted GDF15 and subsequently circulating GDF15 concentrations while also suppressing the cachexia phenotype. Overall, these findings link *STK11/LKB1*-mutated NSCLC cachexia with the cachexia factor GDF15. Of clinical importance, these findings also indicate the need to test patients with treatment naïve *STK11/LKB1*-mutated tumors to determine if they might benefit from anti-GDF15 therapies to suppress wasting due to their tumor-intrinsic biology, independent of therapy-induced cachexia.

## RESULTS

### Characterization of GDF15 tumor and circulating levels across a NSCLC cachexia cohort

We previously identified 17 human non-small lung cancer (NSCLC) lines that produced an unambiguous phenotype of cachexia (n=10) or non-cachexia (n=7) when injected into immunodeficient mice.^1^ Of the 10 lines that induced host wasting, 8 lines had genetic variations in *STK11/LKB1* with moderate to high predicted functional impact. In a candidate approach to identify GDF15’s relevance to STK11/LKB1-mediated cachexia in NSCLC, we assessed tumor *GDF15* RNA expression and systemic GDF15 serum concentrations in the context of host wasting in the cachexia (*red*) and non-cachexia (*black*) lines from this cohort (**Figure 1A**). *GDF15* gene expression was assessed by quantitative reverse transcriptase PCR using human (tumor cell-derived) and mouse (host cell-derived) primers to distinguish tumor cell and host cell changes, respectively. Expression of human *GDF15* mRNA was significantly elevated in a majority of the CC lines compared to non-cachexia lines (**Figure 1B**), with the exception of the H1395 and H1993 cachexia lines that had comparable levels to the non-cachexia cohort. Mouse *Gdf15* mRNA expression in tumors, representing host cell contribution to tumor expression, was statistically unchanged between the cachexia and non-cachexia groups (**Figure 1C**).

**Figure 1.**
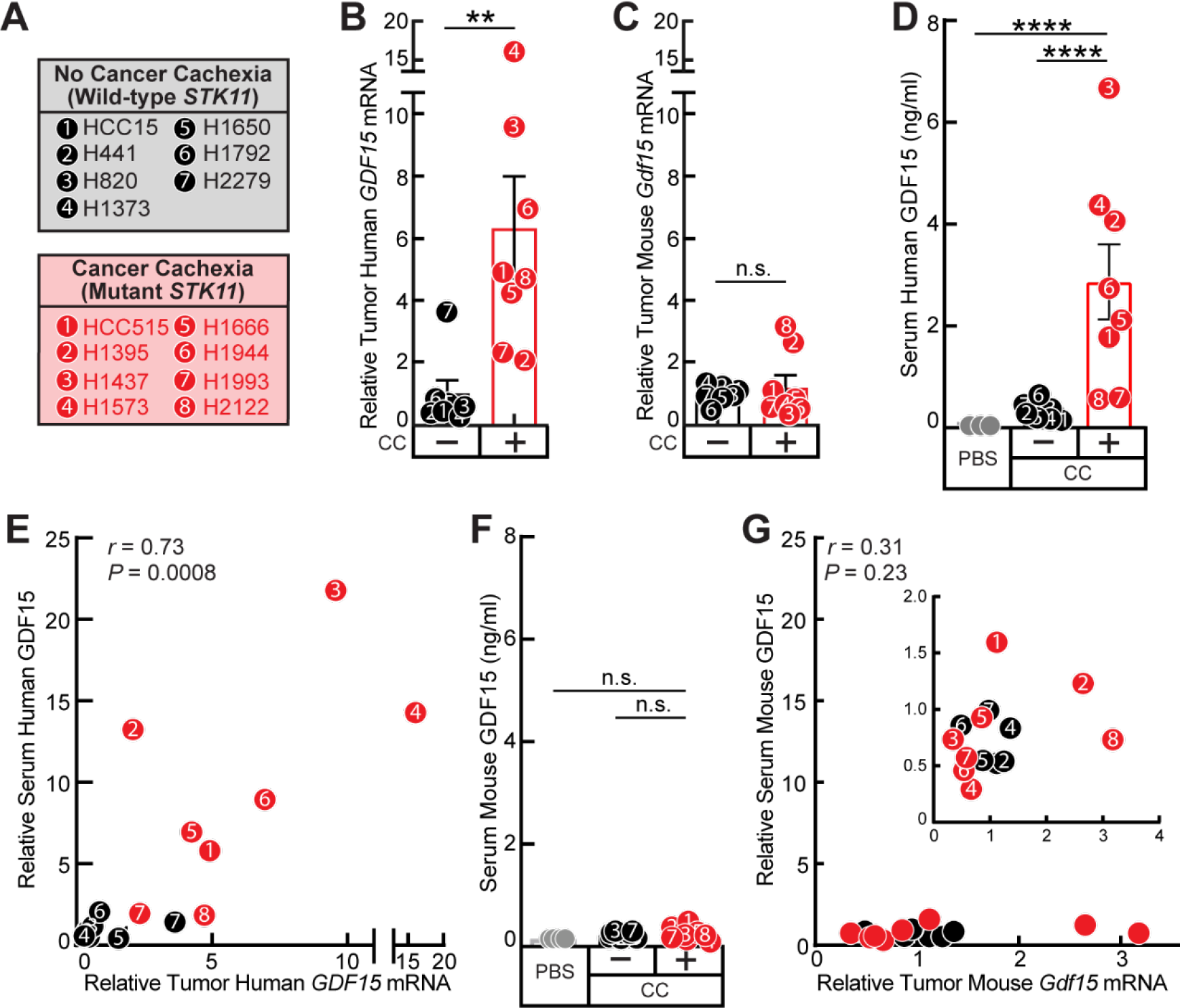
Levels of mRNA and circulating GDF15 of human NSCLC cachexia. A) List of non-cachexia NSCLC lines with wild-type *STK11/LKB1* and cachexia NSCLC lines with mutant *STK11/LKB1* from which tumors and serum have been obtained from previous work when lines were allotransplanted *in vivo*.^1^ B-G) Tumors were harvested and subjected to quantitative RT-PCR for human GDF15 (B,E) and mouse (C,G) *Gdf15* mRNA normalized to β-actin. Serum was subjected to the human (D,E) or mouse (F,G) GDF15 ELISA as described in Methods. Data are shown as mean ± SEM (B-D,F) or as a scatter plot (E,G) relative to the average non-cachexia cohort (B-C,E,G) or the actual measurements (D-G). P was calculated based on unpaired 2-tailed *t* test (B and C) or 1-way ANOVA followed by Dunnett’s multiple-comparison test (D and F), or from linear regression of *GDF15* mRNA levels with serum GDF15 level (E and G).

To evaluate serum concentrations of GDF15 from the mice transplanted with the human cancer lines, we first established human (**Figure S1A**) and mouse (**Figure S1B**) specific GDF15 ELISAs that only recognized human GDF15 (tumor cell-intrinsic source) and murine GDF15 (host cell source), respectively. Serum concentrations of human GDF15 were significantly elevated in mice injected with the cancer cachexia-inducing *STK11/LKB1*-mutated NSCLC lines compared to the non-cachexia wild-type STK11 NSCLC lines and the PBS control cohort (**Figure 1D**), whereas the serum mouse GDF15 concentration was statistically unchanged between the cachexia, non-cachexia, and PBS control groups (**Figure 1F**). Importantly, there was a strong correlation between tumor human *GDF15* mRNA expression and serum human GDF15 concentrations (**Figure 1E**). By contrast, there was no significant association of tumor mouse *Gdf15* mRNA expression with serum mouse GDF15 concentrations (**Figure 1G**). Overall, this data suggested that loss of functional STK11/LKB1-driven NSCLC cachexia is correlated with increases in *GDF15* mRNA expression and serum concentrations that are derived from the tumor cells and not the host.

### Human STK11/LKB1-mutant NSCLC dependence on high circulating GDF15 concentrations for cachexia induction

We next sought to identify if NSCLCs capable of inducing cachexia secondary to *STK11/LKB1* loss are dependent on GDF15 to induce wasting. To test this hypothesis, we used an *STK11/LKB1* mutant, CC-inducing patient line H1573 that exhibited the highest tumor *GDF15* mRNA expression (see **Figure 1B**) and the second highest serum concentration of tumor-derived GDF15 (see **Figure 1D**) when injected into immunocompromised mice. We treated H1573 tumor-bearing mice with either PBS, non-specific IgG monoclonal antibody, or a neutralizing antibody against GDF15 (**Figure 2**). Administration of GDF15 neutralizing antibody or control IgG antibody had no effect on H1573 tumor growth compared to PBS-treated tumor-bearing mice (**Figure 2A**). Human ELISA analysis of serum demonstrated that the administration of GDF15 antibody significantly neutralized the serum human GDF15 concentrations from ∼5 ng/ml in the PBS- and IgG-treated mouse cohorts to ∼0 ng/ml in mice (**Figure 2B**). Mice xenotransplanted with H1573 and administered GDF15 neutralizing antibody had a preservation of body weight (**Figure 2C**) without loss of fat (**Figure 2D**) or lean body mass (**Figure 2E**) compared to PBS- and IgG-treated mice. Thus, treatment with GDF15 neutralizing antibody dramatically suppressed the development of cachexia-associated fat/lean mass and body weight loss normally induced by the *STK11/LKB1*-mutant H1573 NSCLC line.

**Figure 2.**
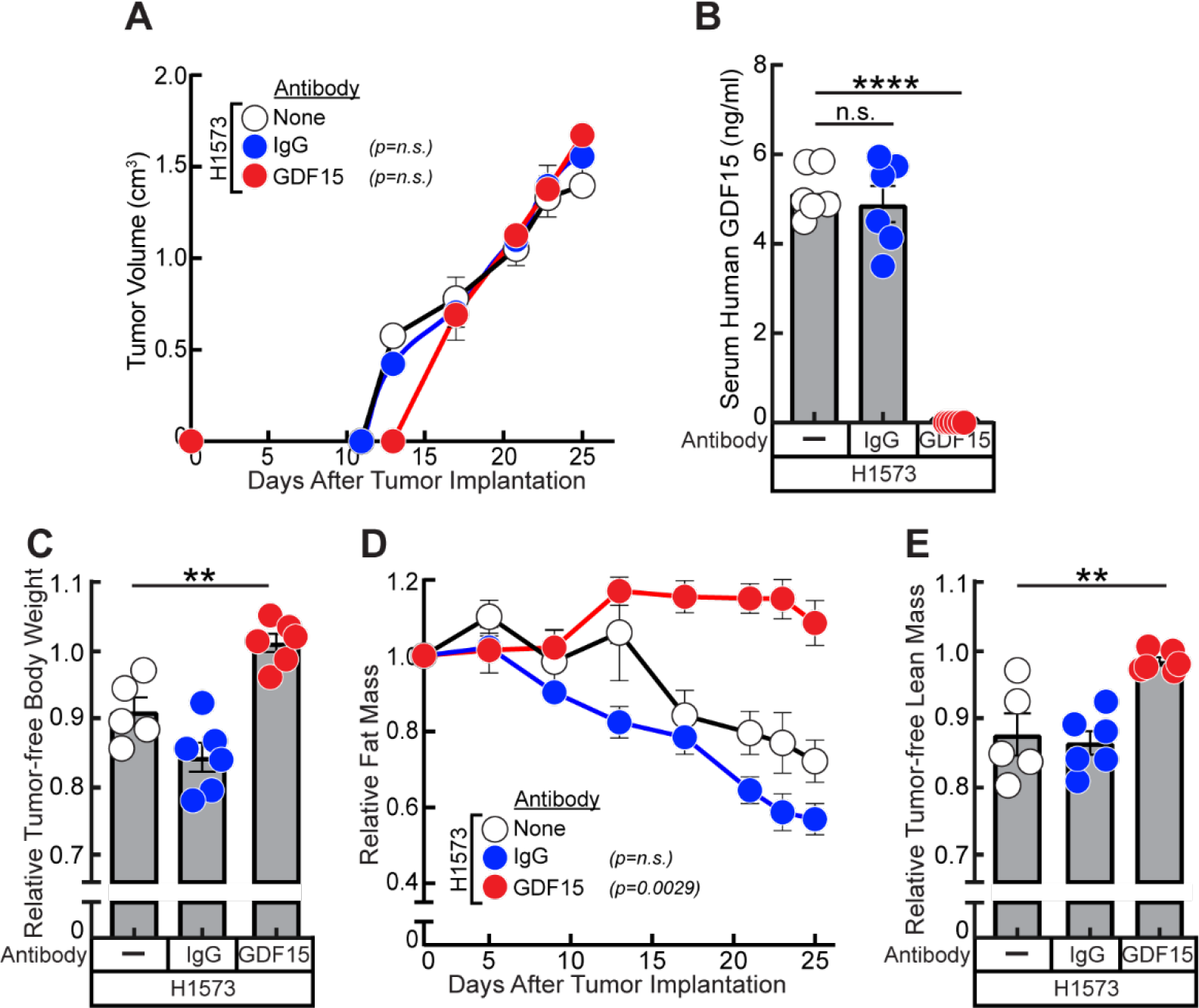
GDF15 antibody neutralization in mice bearing *STK11/LKB1*-mutated NSCLC tumors with high circulating GDF15 concentrations. A-E) Chow-fed 12-13-week-old NOD/SCID male mice (n = 6) were injected s.c. with 200 μl PBS containing 1 x 10^7^ cells from parental H1573 human NSCLC line on day 0. Starting on day 7, mice were injected with s.c. 150 μl PBS in the absence or presence of 10 mg/kg IgG or GDF15 antibody weekly. Longitudinal or endpoint measurements of tumor volume (A), tumor-free body weight (C), fat mass (D), and tumor-free lean mass (E) were obtained as described in Methods. Serum at sacrifice was processed for human GDF15 ELISA (B) as described in Methods. Data are shown as mean ± SEM of the actual measurements (A-B) or relative to their day 0 values (C-E). P was calculated using 1-way (B, C, E) or 2-way (A and D) ANOVA followed by Dunnett’s multiple-comparison test for significant differences from the H1573 parental cohort. *\*\*p<0.01; ***p<0.001; p**** <0.0001; n.s. = not significant; M = GDF15 mature protein; P = GDF15 proprotein*.

The antibody against GDF15 used in our studies is reported to neutralize both human and murine orthologs.^21^ To specifically establish that tumor cell-secreted GDF15 initiates cachexia induction, the H1573 NSCLC cell line was infected with lentiviral CRISPR/Cas9 constructs containing guide RNA targeting *GDF15* (H1573^ΔGDF15^) to silence expression. As a control, the parental H1573 line was also infected with guide RNA directed against *GFP* (H1573^ΔGFP^). Immunoblot analysis of the engineered H1573 cells determined that *GDF15* was only silenced in the H1573^ΔGDF15^ tumor cells compared to the parental H1573 and the H1573^ΔGFP^ tumor cells (**Figure 3A**). These three lines and a PBS vehicle control were subsequently injected subcutaneously into NOD/SCID mice followed by measurement of tumor growth, food intake, body composition, and body weight over time. As shown in **Figure 3B**, tumors derived from the parental H1573, H1573^ΔGFP^, and H1573^ΔGDF15^ cell lines showed similar tumor growth kinetics. Immunoblot analysis revealed that tumors derived from the H1573^ΔGDF15^ line were durably suppressed from expressing GDF15 (**Figure 3C**). Compared to mice harboring parental H1573 or H1573^ΔGFP^ tumors demonstrating serum GDF15 concentrations of ∼4 ng/ml, serum from mice xenotransplanted with H1573^ΔGDF15^ tumors had undetectable human (tumor-derived) GDF15 concentrations to levels comparable to findings in non-tumor bearing PBS-injected mice (**Figure 3D**). ELISA measurements also revealed no significant changes in the serum mouse (host cell-derived) GDF15 concentrations in parental or engineered lines compared to the PBS control (**Figure 3E**), suggesting a lack of host contribution to serum GDF15 levels. Silencing of *GDF15* did not cause a significant increase in food intake in mice compared to the mice injected with parental H1573 or H1573^ΔGFP^ cells (**Figure 3F**). Critically, compared to the mice injected with parental H1573 or H1573^ΔGFP^ cells, mice injected with the H1573^ΔGDF15^ cell line had a rescue of their body weight (**Figure 3G**), tumor-free body weight (**Figure 3H**), lean mass (**Figure 3I**), tumor-free lean mass (**Figure 3J**), and fat mass (**Figure 3K**), similar to levels observed in the PBS-injected, non-tumor bearing mice. Preservation of host adipose tissue with tumor cell *GDF15* silencing was further verified by H&E staining of epididymal adipose tissue (**Figure 4A**) and by measurement of adipocyte cross-sectional area which favored larger area adipocytes compared to those adipocytes taken from mice harboring parental H1573 or H1573^ΔGFP^ tumors (**Figures 4B-C**). Concordant with the measurements of lean mass by ECHO-MRI, metachromatic ATPase staining of the gastrocnemius muscle ATPase activity revealed a significant increase of the total and type 1 muscle fiber cross-sectional area of the mice with the H1573^ΔGDF15^ tumors compared to mice harboring the parental or H1573^ΔGFP^ tumors (**Figures 4D-E**). Although the mice containing H1573^ΔGDF15^ tumors showed a significant increase of type 2 muscle fiber cross-sectional area when compared to the parental H1573 line, this increase was not significant when compared to mice with H1573^ΔGFP^ tumors (**Figure 4E**). Overall, these studies proved that *STK11/LKB1*-mutated H1573 NSCLCs intrinsically rely on tumor cell-derived GDF15 for promotion of cachexia-associated adipose and muscle atrophy.

**Figure 3.**
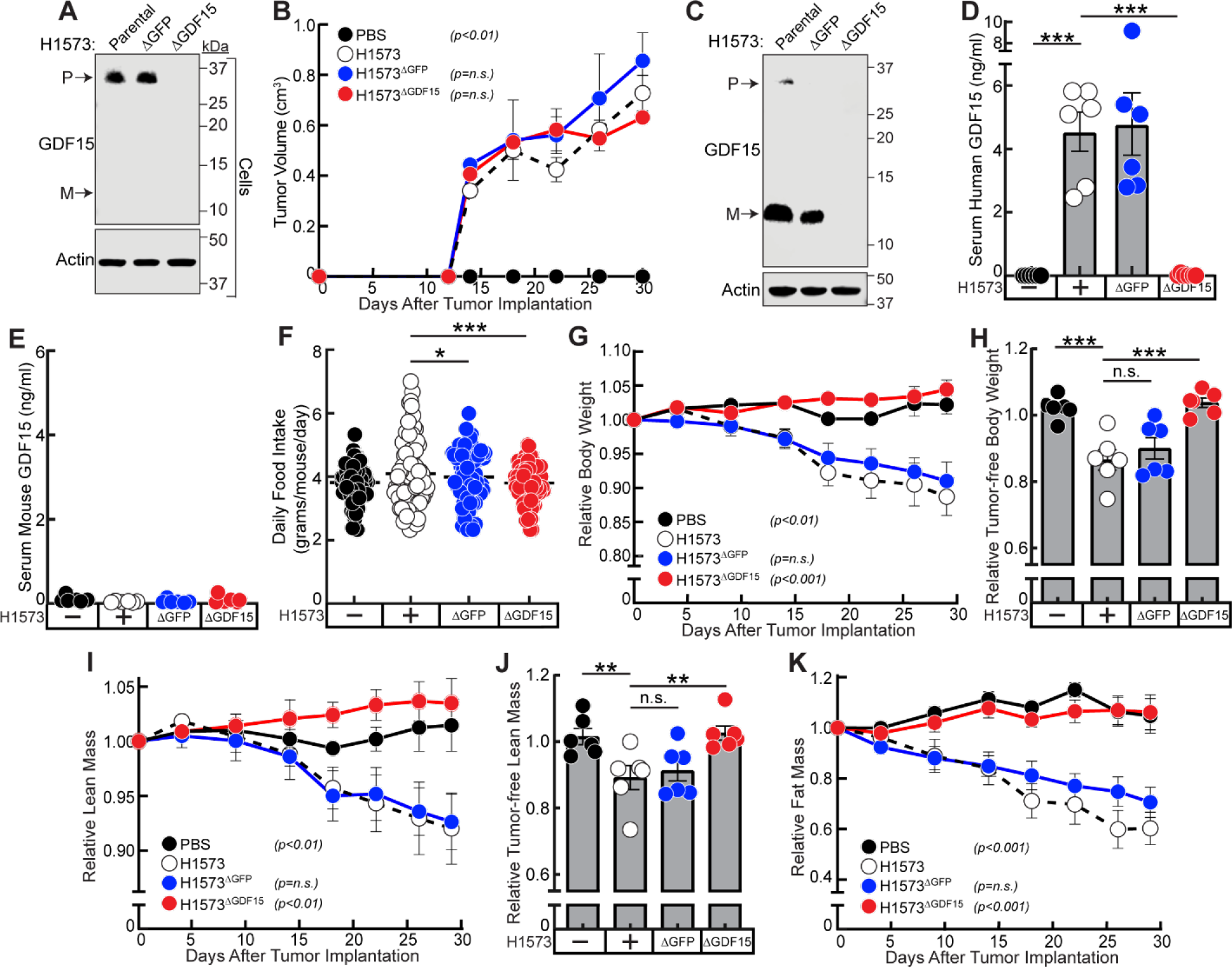
*GDF15* silencing in *STK11/LKB1*-mutated NSCLC tumors allotransplanted into mice that have high circulating GDF15 concentrations. A-K) Chow-fed 13-week-old NOD/SCID male mice (n = 6 per group) were injected s.c. with 200 ul PBS in the absence or presence of 1 x 10^7^ cells from parental H1573, H1573^ΔGFP^, or H1573^ΔGDF15^ lines as described in Methods. Longitudinal or endpoint measurements of tumor volume (B), food intake (F), body weight (G), tumor-free body weight (H), lean mass (I), tumor-free lean mass (J), or fat mass (K) were obtained as described in Methods. Tumor cells before injection (A) and tumors at sacrifice (C) were processed for immunoblot analysis with the indicated antibody. Serum at sacrifice was subjected to the human (D) or mouse (E) GDF15 ELISA as described in Methods. Data are shown as mean ± SEM of the actual measurements (B, D-F) or relative to their day 0 values (G-K). P was calculated using 1-way (D-F, H, J) or 2-way (B, G, I, K) ANOVA followed by Dunnett’s multiple-comparison test for significant differences from the H1573 parental cohort. *\*p<0.05; **p<0.01; p***<0.001; n.s. = not significant; M= GDF15 mature protein; P = GDF15 proprotein*.

**Figure 4.**
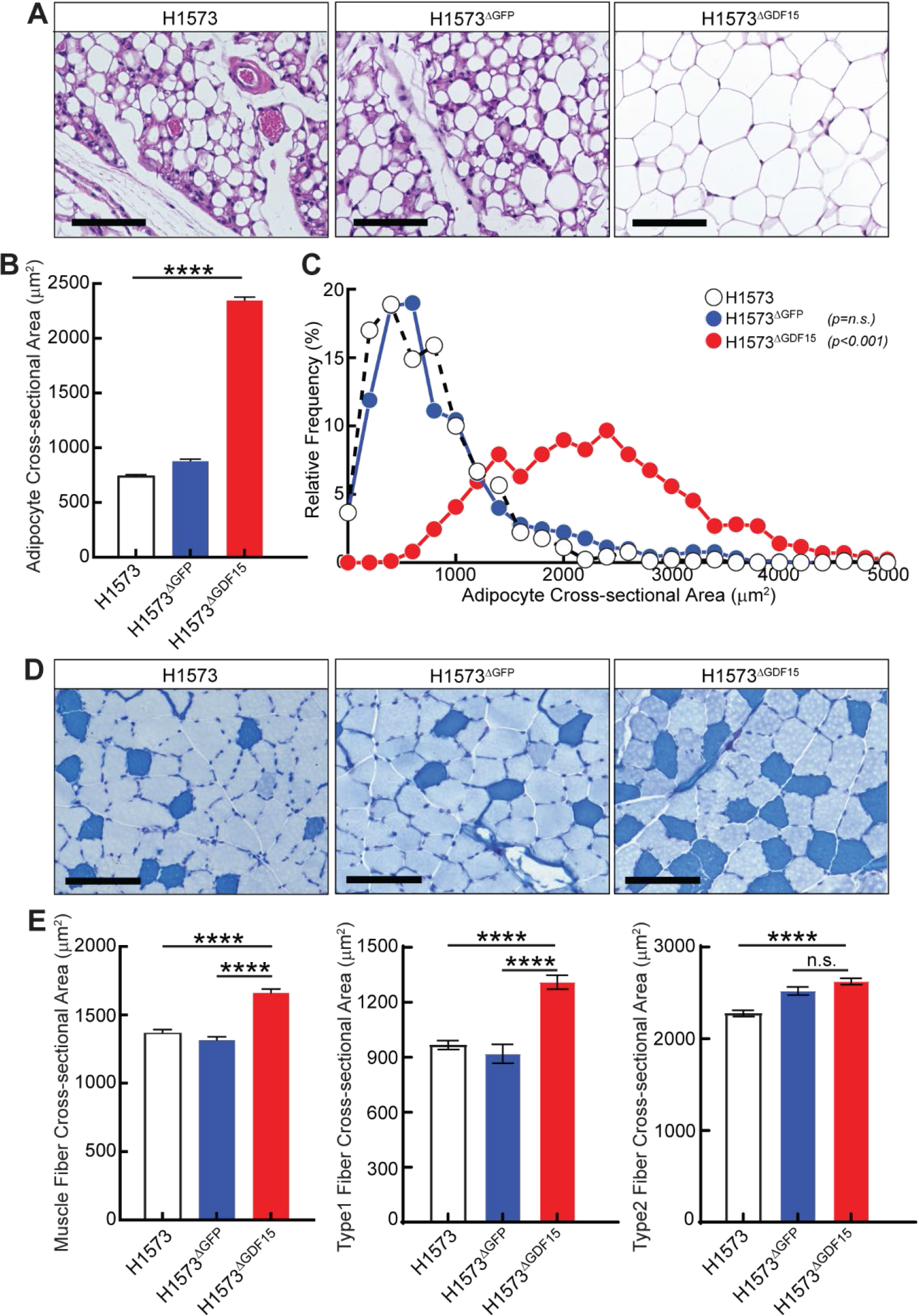
Analysis of adipose and muscle obtained from isogenic H1573 NSCLC lines allotransplanted into murine models. A-E) NOD/SCID mice were treated as described in Figure 3. Mice were sacrificed with epididymal white adipose tissue (eWAT) and gastrocnemius muscle harvested and processed for H&E (A) or metachromatic ATPase staining (D) staining, respectively (scale bar 100 μm). Subsequently, measurements of adipocyte (B-C) and myocyte (E) cross-sectional area from 3 mice per cohort were obtained. P was calculated using 1-way (B, C, E) followed by Dunnett’s multiple-comparison test for significant differences from the H1573 parental cohort. *p****<0.0001; n.s. = not significant*

### Human STK11/LKB1-mutant NSCLC dependence on intermediate circulating GDF15 concentrations for cachexia induction

Having established that mice xenotransplanted with human NSCLC lines demonstrating high circulating human GDF15 concentrations (∼4 ng/ml or more) are likely dependent on tumor-derived GDF15 for cachexia induction, we next sought to determine if intermediate serum concentrations of human GDF15 (∼2 ng/ml) were sufficient to support cachexia-associated adipose and muscle wasting induced by NSCLCs possessing *STK11/LKB1* loss of function variants. The parental human 1944 NSCLC line had average human *GDF15* expression (see **Figure 1B**) and intermediate serum concentrations (∼2ng/ml; see **Figure 1D**) compared to the other *STK11*/*LKB1* mutant CC inducing lines. We therefore separately silenced *GDF15* (H1944^ΔGDF15^) or *GFP* (H1944^ΔGFP^) in the parental H1944 cells using lentiviral CRISPR/Cas9 techniques. Immunoblot analysis verified that that the monoclonal H1944^ΔGDF15^ line had silenced GDF15 protein expression compared to parental H1944 and H1944^ΔGFP^ control cell lines (**Figure S2A**). These parental and engineered cell lines were subsequently injected subcutaneously into NOD/SCID immunodeficient mice followed by longitudinal evaluations of tumor growth, food intake body weight, lean mass, and fat mass. Although tumors derived from the H1944^ΔGDF15^ cells showed persistent elimination of GDF15 protein expression by immunoblot analysis (**Figure S2A**), there was no significant change in tumor growth when comparing the parental H1944, H1944^ΔGFP^, and H1944^ΔGDF15^ tumors (**Figure S2B**). Serum from mice harboring H1944^ΔGDF15^ tumors had a significant decrease in human (tumor cell-derived) GDF15 concentration compared to the serum of mice injected with parental H1944 or H1944^ΔGFP^ cells (**Figure S2C**). However, serum mouse (host cell-derived) GDF15 concentrations remained low in all three cohorts (**Figure S2D**). In this model, mice harboring H1944^ΔGDF15^ tumors had a significant increase in their daily food intake compared to the control cohorts (**Figure S2E**). Furthermore, compared to the mice injected with parental H1944 or H1944^ΔGFP^ cells, mice injected with the H1944^ΔGDF15^ cell line also exhibited a rescue of their tumor-free body weight (**Figure S2F**), tumor-free lean mass (**Figure S2G**), and fat mass (**Figure S2H**). These studies confirmed that intermediate concentrations of circulating GDF15 derived from an *STK11/LKB1*-mutated NSCLC line can also induce the cachexia phenotype and is rescued by functional genomic targeting of tumor cell *GDF15* expression.

### Human STK11/LKB1-mutant NSCLC dependence on low circulating GDF15 concentrations for cachexia induction

Using GDF15 neutralizing antibody and/or tumor cell *GDF15* silencing studies, we were able to clearly show that cancer cachexia models driven by *STK11/LKB1* loss of function tumor variants with high or even intermediate concentrations of circulating GDF15 were dependent on this tumor-derived factor to promote cachexia-associated fat and muscle atrophy. We used the next set of studies to establish a lower GDF15 concentration boundary of dependence by *STK11/LKB1*-mutated NSCLC lines for promoting cachexia wasting. Although immunodeficient mice harboring *STK11/LKB1*-mutated H2122 tumors induce cachexia when injected subcutaneously into NOD/SCID mice, their human (tumor-derived) *GDF15* mRNA expression (see **Figure 1B**) and GDF15 serum concentrations (see **Figure 1D**) were at levels similar to mice bearing NSCLCs with wild-type *STK11/LKB1* that could not induce cachexia. In **Figure S3**, we administered immunodeficient mice xenotransplanted with H2122 cells either PBS, non-specific IgG monoclonal antibody, or a neutralizing antibody against GDF15. Administration of GDF15 neutralizing antibody or control IgG antibody had no effect on H2122 tumor growth compared to PBS-injected tumor-bearing mice (**Figure S3A**). Serum GDF15 ELISA evaluations demonstrated that the administration of GDF15 neutralizing antibody significantly decreased the serum human GDF15 concentrations from the PBS- and IgG-treated mouse cohorts to background levels in mice administered GDF15 neutralizing antibody (**Figure S3B**). Although GDF15 neutralizing antibody administered to H2122-bearing mice did not significantly affect food intake (**Figure S3C**), there was a significant preservation of fat mass (**Figure S3E**). However, this suppression of fat mass loss was not complete and durable since GDF15 antibody-treated mice still lost ∼20% of their fat mass compared to their Day 0 values. There was a trend for body weight preservation in the H2122-bearing mice treated with GDF15 antibody compared to control cohorts, but this was not statistically significant (**Figure S3D**).

Since the H2122 cachexia phenotype was partially responsive to GDF15 neutralizing antibody, we next silenced *GDF15* (H2122^ΔGDF15^) or *GFP* (H2122^ΔGFP^) in the parental H2122 cells using lentiviral CRISPR/Cas9 techniques. Immunoblot analysis revealed that the H2122^ΔGDF15^ line had suppression of GDF15 protein expression compared to parental H2122 and H2122^ΔGFP^ cell lines (**Figure 5A**). These cell lines were subsequently injected subcutaneously into NOD/SCID immunodeficient mice followed by measurement of the wasting phenotype. There was no significant difference in tumor growth among the three tumor cohorts (**Figure 5B**) despite the persistent silencing of GDF15 protein expression from the H2122^ΔGDF15^ tumors (**Figure 5A**). Mice harboring the parental H2122 or the H2122^ΔGFP^ lines had average serum human (tumor cell-derived) GDF15 concentrations less than 0.5 ng/ml (**Figure 5C**). Although this was significantly increased compared to PBS-treated mice, this concentration of GDF15 was an order of magnitude lower than observed in the H1573 (see **Figure 3D**) and H1944 (see **Figure S2C**) mouse studies. As expected, mice harboring H2122^ΔGDF15^ tumors had a significant decrease in human (tumor cell-derived) GDF15 concentrations (**Figure 5C**). With respect to resulting phenotypes, we did observe a small but significant increase in food intake in the mice harboring H2122^ΔGDF15^ tumors compared to those mice bearing parental tumors (**Figure 5E**). There was also a significant preservation of body weight (**Figure 5F**), tumor-free lean mass (**Figure 5G**), and fat mass (**Figure 5I**) compared to the parental H2122- and H2122^ΔGFP^-bearing mice. In concordance, there was an increase in average myocyte (**Figure 5H**) and adipocyte (**Figure 5J**) cross-sectional area in mice transplanted with H2122^ΔGDF15^ tumors compared to mice harboring parental H2122 or H2122^ΔGFP^ tumors. Overall, tumor-secreted GDF15 also contributed to the cachexia phenotype in mice bearing *STK11*/LKB1-mutated NSCLC lines demonstrating low systemic concentrations of the protein.

**Figure 5.**
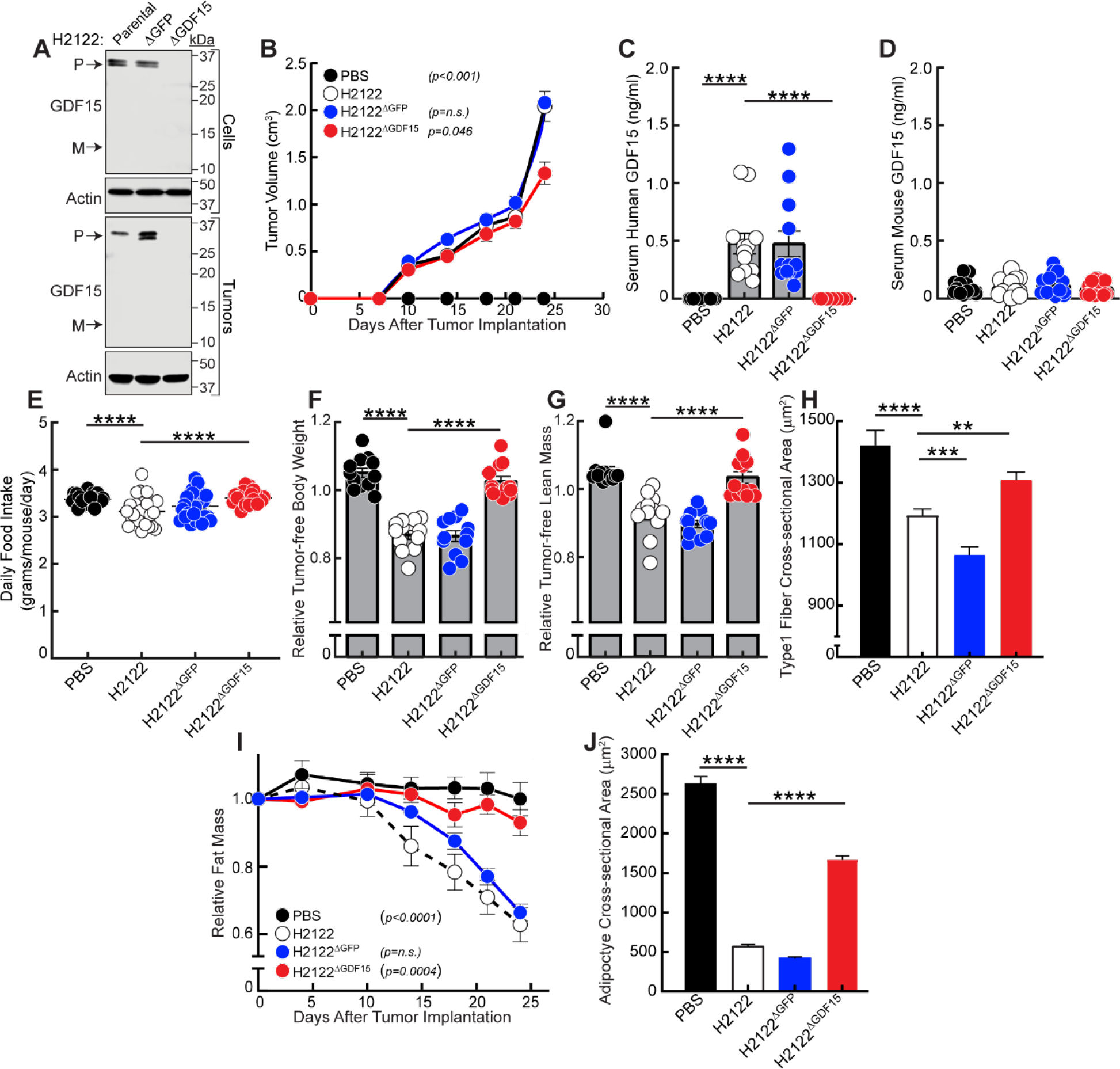
*GDF15* silencing in *STK11/LKB1*-mutated NSCLC tumors allotransplanted into mice that have low circulating GDF15 concentrations. A-J) Chow-fed 13-week-old NOD/SCID male mice (n = 12 per group, combination of two experiments) were injected s.c. with 200 ul PBS in the absence or presence of 1 x 10^7^ cells from parental H2122, H2122^ΔGFP^, or H2122^ΔGDF15^ lines as described in Methods. Longitudinal or endpoint measurements of tumor volume (B), food intake (E), tumor-free body weight (F), tumor-free lean mass (G), or fat mass (I) were obtained as described in Methods. Tumor cells before injection and tumors at sacrifice were processed for immunoblot analysis with the indicated antibody (A). At sacrifice, serum was subjected to the human (C) or mouse (D) GDF15 ELISA and gastrocnemius muscle and epididymal white adipose tissue were harvested and myocyte and adipocyte cross sectional area was measured as described in Methods. Data are shown as mean ± SEM of the actual measurements (B-E, H, J) or relative to their day 0 values (F-H). P was calculated using 1-way (C-H, J) or 2-way (B, I) ANOVA followed by Dunnett’s multiple-comparison test for significant differences from the H2122 parental cohort. *p****<0.0001; n.s. = not significant; M = GDF15 mature protein; P = GDF15 proprotein*.

### GDF15 dependence in NSCLC STK11/LKB1 isogenic model systems of cachexia

To this point, we had focused on studying cachexia-inducing human NSCLCs with endogenous loss of *STK11/LKB1* from spontaneous mutations. These tumors, however, do possess co-occurring mutations and other alterations potentially affecting or contributing to cachexia development. Furthermore, we wanted to develop an isogenic system permitting us to better understand how contributory changes in only STK11/LKB1 function were to cachexia development *in vivo*. Previously, we silenced *STK11*/*LKB1* (H1792^ΔSTK11^) in the parental H1792 NSCLC line to convert a non-cachexia inducing tumor line into one with cachexia-inducing potential without changing the mRNA expression of human (tumor cell-derived) *GDF15*.^1^ To test if this line was also dependent on GDF15 to induce host wasting, we injected immunocompromised mice with H1792^ΔSTK11^ cells followed by administration of control IgG or anti-GDF15 monoclonal antibodies. In **Figure S4A**, we established that that the H1792^ΔSTK11^ line maintained similar tumor growth kinetics even when xenotransplanted into mice treated with either GDF15 antibody or non-specific IgG antibody. In **Figure S4B**, we verified that mice bearing H1792^ΔSTK11^ tumors had similar human GDF15 serum concentrations below the detection limit of the assay to serum obtained from mice bearing control 1792 lines. Therefore, we used this model as another potential validation of GDF15’s role in cachexia induction even at very low serum concentrations. Again, systemic anti-GDF15 antibody compared to control IgG treatment reduced the potential of H1792^ΔSTK11^ tumors to cause weight loss (**Figure S5C**), lean mass loss (**Figure S5D**), and adipose loss (**Figure S5E**) *in vivo*. From the collective data from the above models, GDF15 contributes to the cachexia phenotype in *STK11/LKB1*-mutant NSCLC across a wide range of serum concentrations and independent of iatrogenic effects of cytotoxic chemotherapies.

### Dependence of STK11/LKB1-mutated NSCLC cachexia on GDF15 is verified in syngeneic models

Previously, all experiments had been conducted using human NSCLCs derived from patients in immunodeficient murine models. To determine if similar findings could be recapitulated with an intact immune system, we obtained murine isogenic tumors cell lines derived from well-established KP and KPL genetically-engineered mouse models (GEMMs).^22^ The KP model has mutations in *Trp53* and *Kras* leading to functional inactivation of both genes whereas the KPL model has an additional inactivating mutation in *Stk11/Lkb1*. Use of tumor cells derived from these murine lines in allotransplant experiments permitted us the ability to titrate the timing of tumor generation and kinetics of tumor growth in the context of a fully functional immune system. As expected, the KPL cell line expressed no STK11/LKB1 and had limited phosphorylation of AMPK (**Figure 6A**) compared to the KP line by immunoblot analysis. Furthermore, the allotransplant lines had similar tumor growth kinetics (**Figure 6B**) in their syngeneic hosts. Food intake (**Figure 6C**) was reduced in the mice bearing KPL tumors compared to controls. Unlike tumors formed from the KP line, however, those formed from transplantation of the KPL line were able to induce cachexia-associated tumor-free body weight loss (**Figure 6E**), tumor-free lean mass loss (**Figure 6F**), and adipose loss (**Figure 6G**). Although there was ∼2-fold increase in serum mouse GDF15 levels in KPL-bearing mice compared to KP-bearing mice, these serum concentrations were lower than all of the patient-derived STK11/LKB1-mutated NSCLC lines previously tested (see **Figure 1D**). To determine if GDF15 is relevant to cachexia induced in KPL-bearing animals, we suppressed GDF15 action with administration of GDF15 neutralizing antibody to specific KPL-bearing mouse cohorts. These studies verified that KPL-induced wasting metrics – anorexia (**Figure 6I**), adipose loss (**Figure 6J**), and lean mass loss (**Figure 6K**) – were dependent on GDF15 despite similar tumor growth (**Figure 6H**). Therefore, this syngeneic model further validated GDF15’s role in STK11/LKB1-mutated cachexia induction even at very low serum concentrations.

**Figure 6.**
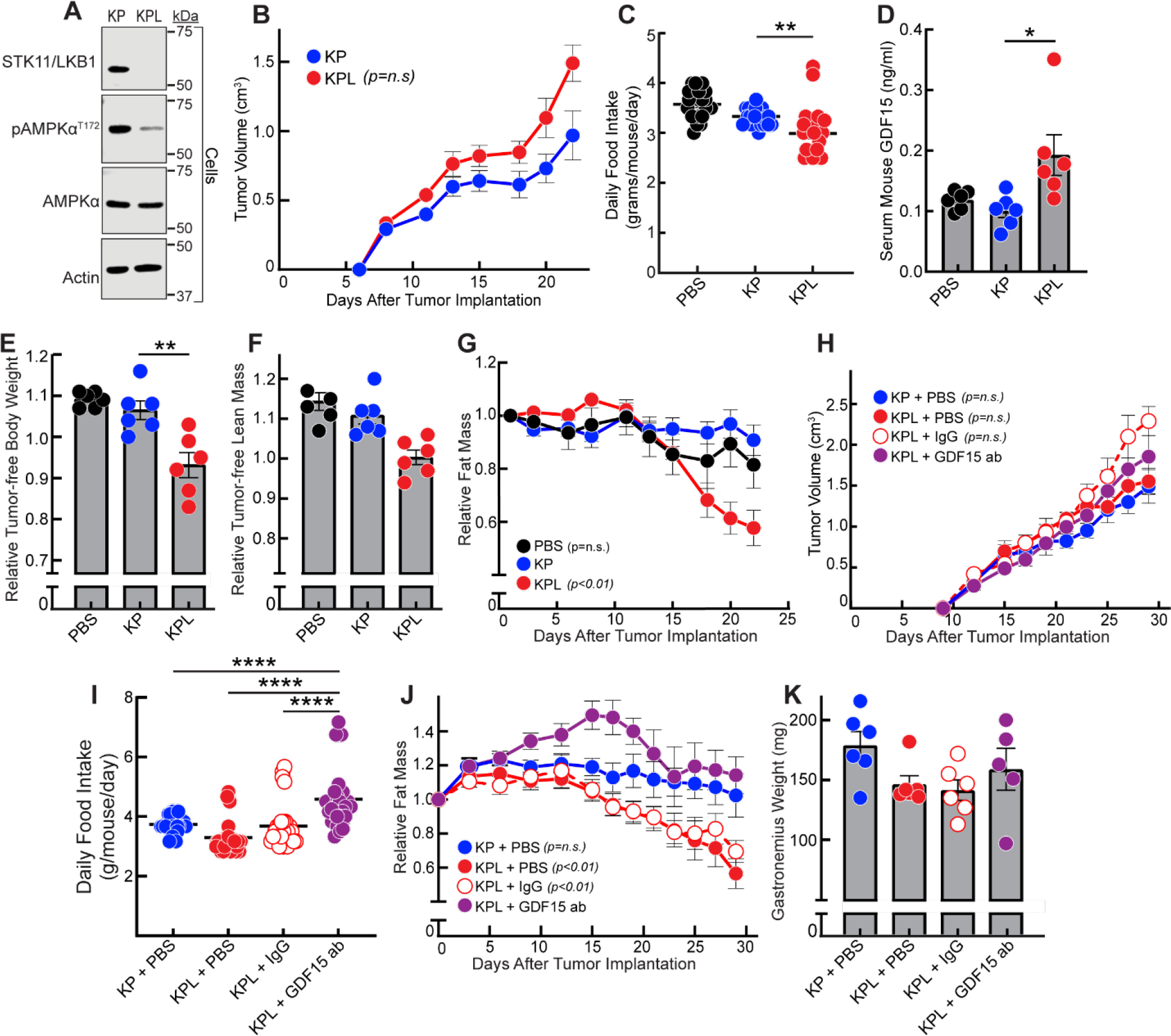
Cachexia Phenotype and GDF15 antibody neutralization of syngeneic mice bearing KP and KPL tumors. A) Tumor cells before injection were processed for immunoblot analysis with the indicated antibodies. B-K) Chow-fed 12-week-old C57BL6J male mice (n = 6) were injected s.c. with 200 μl PBS containing in the absence or presence of 1 x 10^6^ cells from the mouse NSCLC KP or KPL lines on day 0 (A-G) followed by weekly injections of 150 μl PBS in the absence or presence of 10 mg/kg IgG or GDF15 antibody (H-K) Longitudinal or endpoint measurements of tumor volume (B,H), food intake (C,I), tumor-free body weight (E), tumor-free lean mass (F), and fat mass (G,J) were obtained as described in Methods. At sacrifice, serum was processed for mouse GDF15 ELISA (D) as described in methods and the gastrocnemius was harvested and weighed. Data are shown as mean ± SEM of the actual measurements (B-D,H-I,K) or relative to their day 0 values (E-G,J). P was calculated using 1-way (C-F,I,K) or 2-way (B,G,H,K) ANOVA followed by Dunnett’s multiple-comparison test for significant differences from the KP (B-G) or the KPL + GDF15 antibody (H-K) cohorts. *p* <0.05; p** <0.01, p**** <0.0001; n.s. = not significant*.

### Reconstitution of wild-type STK11/LKB1 verifies its role in regulating cachexia through regulation of GDF15 expression and release

Tumor-bearing parental H1437 mice had the second highest tumor mRNA expression and the highest serum concentrations of human tumor-cell derived GDF15 when compared to the other *STK11/LKB1*-mutant lines and the non-cachexia lines as well (see **Figures 1B and 1D**). To assess if we could abrogate cachexia induction in these *STK11/LKB1*-mutant NSCLC lines with reconstitution of wild-type STK11/LKB1 protein, we overexpressed the wild-type gene using a lentivirus system and made stable H1437^STK11^ tumor lines. We also created a monoclonal control line that received only empty vector (H1437^CTL^). As observed from immunoblotting analysis in **Figure 7A**, we were successfully able to create two monoclonal lines overexpressing wild-type STK11/LKB1 (H1437^STK11^), associated with the expected phosphorylation of AMPK. This finding was verified in the stable H1437^STK11^ lines created in addition to the tumors derived from these lines *in vivo*. Furthermore, **Figure 7A** also verified that loss of *STK11/LKB1* gene expression and protein in the parental H1437 line led to an expected suppression of phosphorylation and activation of AMPK. Interestingly, reconstitution of wild-type STK11/LKB1 protein blocked induction/expression and processing of GDF15 protein (both pre and mature forms) in cells and tumors (**Figure 7A-B**).

**Figure 7.**
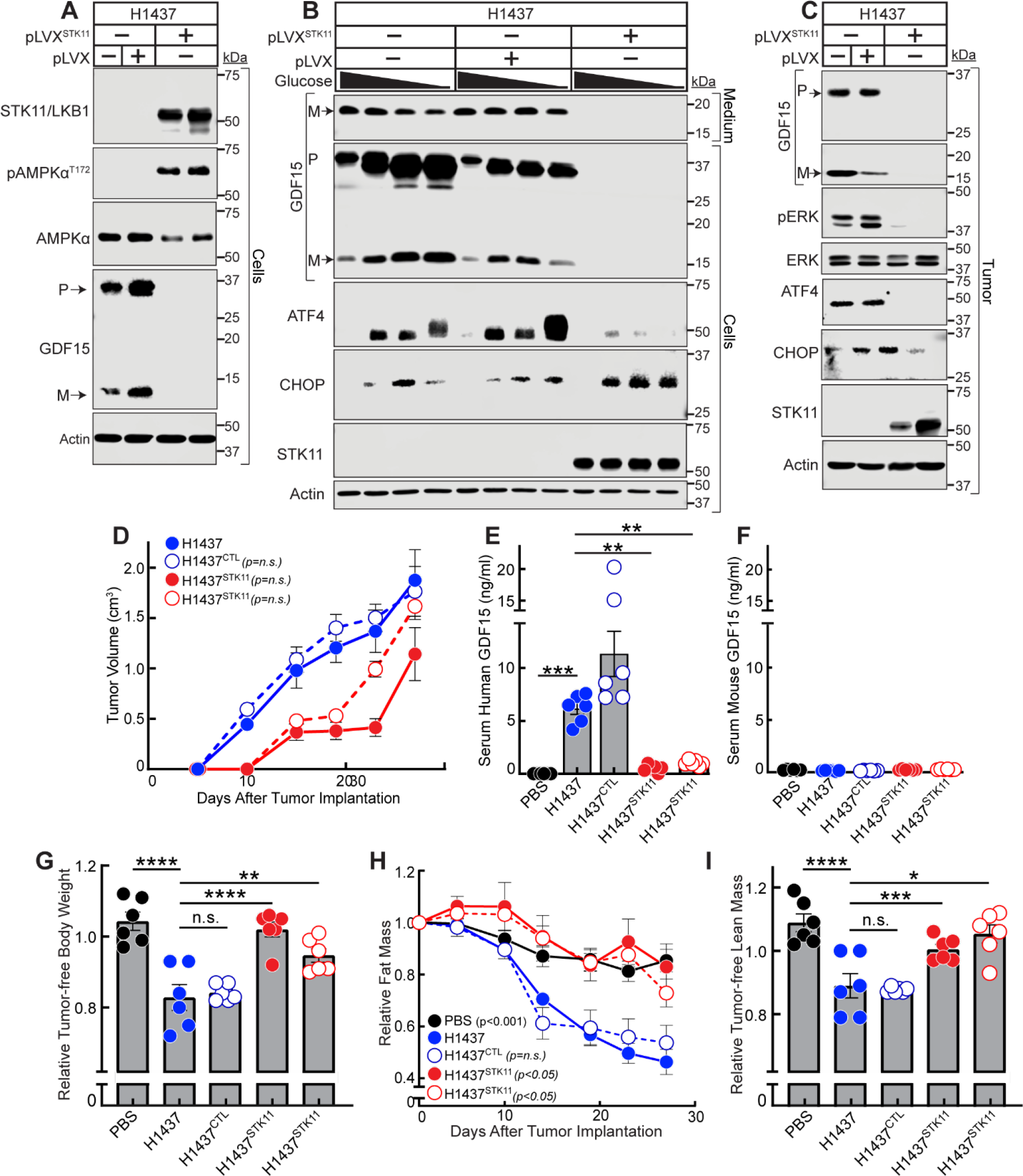
Effect of wild-type STK11/LKB1 reconstitution on GDF15 processing and cachexia induction. A-B) Expression and secretion of GDF15 from patient-derived NSCLC cell line reconstituted with wild-type *STK11/LKB1.* Parental H1437, H1437^CTL^, or H1437^STK11^ cells were set up in medium A at a density of 4 x 10^5^ cells/well of 6-well plates on day 0. On day 1, medium was removed, cells were washed with 2 ml of PBS and harvested (A) or incubated with 2 ml of medium D in the presence of 0.01-10 mM glucose for 16 hr at 37°C followed by harvesting (B). Medium and cell lysate were subjected to immunoblot analysis with the indicated antibody as described in Methods. C-J) Effect of STK11/LKB1 reconstitution on cachexia phenotype. Chow-fed 15-week-old NOD/SCID male mice (n = 6 per group) were injected s.c. with 200 ul PBS in the absence or presence of 1 x 10^7^ cells from parental H1437, H1437^CTL^, H1437^STK11^, or another H1437^STK11^ monoclonal line as described in Methods. Longitudinal or endpoint measurements of tumor volume (D), tumor-free body weight (H), fat mass (I), or tumor-free lean mass (J) were obtained as described in Methods. Tumors at sacrifice were processed for immunoblot analysis with the indicated antibody as described in Methods (A). At sacrifice, serum was subjected to the human (E) or mouse (F) GDF15 ELISA as described in Methods. Data are shown as mean ± SEM of the actual measurements (D, E-F) or relative to day 0 values (G-I). P was calculated using 1-way (E-G, I) or 2-way (D, H) ANOVA followed by Dunnett’s multiple-comparison test for significant differences from the H1437^Ctl^ cohort. *\*p<0.05; **p<0.01; p***<0.001; n.s. = not significant; M= GDF15 mature protein; P = GDF15 proprotein*.

We next tested these cells in conditions of nutrient rich (high glucose) and nutrient deprived (low glucose) *in vitro* medium conditions, especially knowing of the role that GDF15 increases in nutrient-limited states through the activation of ER stress proteins CHOP and ATF4.^23, 24^. In the parental 1437 NSCLC line devoid of wild-type STK11/LKB1 expression, several gene products including GDF15 and the ER stress proteins CHOP and ATF4 were increasingly induced with lowering of glucose/nutrients (**Figure 7B**). When wild-type STK11/LKB1 was reconstituted into the parental 1437 NSCLC line, however, GDF15 expression (pro and mature forms) was completely abrogated and that of ATF4 were nearly so, whereas CHOP expression was similar comparing lines with wild-type and mutant STK11/LKB1. Tumors derived from transplantation of the 1437 line with wild-type STK11/LKB1 reconstituted maintained an expression of the kinase while still suppressing expression of all GDF15 protein forms and additionally phosphorylated forms of ERK as would be expected (**Figure 7C**). These data provided evidence of a clear mechanistic interaction between the presence or absence of STK11/LKB1 protein and the induction and processing of GDF15, the latter necessary for cachexia development *in vivo*. As expected, overexpression of the wild-type version of the tumor suppressor STK11/LKB1 led to a moderate suppression of tumor growth kinetics compared to control tumors (**Figure 7D**), but animals were only sacrificed for cachexia evaluations once all tumors from all cohorts reached similar terminal sizes (**Figure 7D**).

In parallel to the cell and tumor findings, mice bearing the STK11/LKB1 reconstituted H1437^STK11^ tumors demonstrated human serum GDF15 concentrations that approached baseline levels (i.e. equivalent to mice bearing no tumor) in comparison to the elevated serum GDF15 concentrations observed in mice bearing parental 1437 tumors or 1437 tumors overexpressing control vectors (**Figure 7E**). There was no significant change in murine serum GDF15 concentrations among all cohorts (**Figure 7F**). With suppression of GDF15 induction and release into serum, mice bearing H1437^STK11^ tumors displayed an improvement in food intake (**Figure S5A-B**) and preservation of body weight (**Figure 7G**), fat mass (**Figure 7H**), and lean weight (**Figure 7I**) when compared to mice bearing parental 1437 or 1437^ctrl^ tumors. Ultimately, findings from these experiments continue to highlight a dependence of *STK11/LKB1*-mutated NSCLC tumors on GDF15 protein expression and processing for the induction of cachexia wasting.

## DISCUSSION

This study was undertaken to better characterize cachexia induction when non-small cell lung cancers lost STK11/LKB1 functional capacity. Our previous study demonstrated that tumors cells with *STK11/LKB1* loss-of-function variants promoted cachexia associated adipose and muscle loss when transplanted into mice.^1^ Tumor mutant variants in *STK11/LKB1* were also associated with weight loss at cancer diagnosis in advanced NSCLC patients. We, therefore, initiated a candidate approach to identify downstream molecules with metabolic associations that were enriched when STK11/LKB1 function was lost. Given data from other studies, we initially focused on GDF15, a TGFβ family member and anorexia modulator.^3–5^ GDF15 is induced in host tissues of patients in settings of cytotoxic chemotherapy administration or other forms of systemic stress.^9, 10, 13^ Interestingly, previous work has shown that GDF15 expression is regulated in hepatocytes by AMPK, a downstream target of STK11/LKB1 regulation.^25^ We postulated that loss of STK11/LKB1 function in tumor cells could similarly induce GDF15 enrichment locally and systemically in support of wasting. Mice bearing patient-derived human STK11/LKB1 variant NSCLC lines with cachexia-inducing capacity demonstrated elevated serum GDF15 levels compared to mice bearing wild-type STK11/LKB1 NSCLC tumors. Using PCR primers and ELISAs that are capable of distinguishing mouse-versus human-derived GDF15, we identified an enrichment of human serum GDF15 and human tumor GDF15 mRNA in the *STK11/LKB1* variant NSCLC-bearing mice. Use of GDF15 neutralizing antibodies currently being tested in randomized Phase 2 clinical trials suppressed STK11/LKB1-mutated NSCLC cancer-associated adipose and muscle wasting in multiple patient-derived models and isogenic patient-derived and a syngeneic mouse-derived model without affecting tumor progression. To establish that the tumor and not the host was the source of the GDF15 that was relevant to STK11/LKB1 NSCLC cachexia, we silenced *GDF15* transcript tumor-cell specifically and again saw an abrogation of the cachexia development in multiple models demonstrating a spectrum of systemic GDF15 levels, including those with low/undetectable serum GDF15 levels. Finally, we also reconstituted wild-type STK11/LKB11 in a patient-derived NSCLC line containing a STK11/LKB1 gene deletion and reversed GDF15 production and secretion reducing serum levels and cachexia development. In aggregate, these data suggest that tumor-derived GDF15 is a conduit through which STK11/LKB1 loss-of-function in NSCLC supports the development of cachexia-associated adipose, muscle, and body weight loss.

Historically, the field of cachexia has focused on host tissue as being the ultimate source of GDF15-associated anorexia and wasting, particularly under the stress of cytotoxic chemotherapy during tumor-directed therapy.^13^ The association between GDF15 and cachexia is certainly not all that surprising, as multiple groups now have linked cytotoxic chemotherapy to the induction of host GDF15 to promote chemotherapy-associated anorexia/cachexia.^14^ With cachexia incidence still high across multiple primary cancers despite reduced use of cytotoxic chemotherapies, it became apparent that tumor cells themselves could be a contributor to systemic GDF15 levels than previously appreciated. Recently, the Tracer-X study identified an association between serum GDF15 levels and cachexia outcomes.^15^ However, since that group did not identify an association between tumor expression and circulating concentrations of GDF15, they concluded the source of circulating GDF15 was likely derived from the host. Having accumulated 8 patient-derived *STK11/LKB1*-mutant NSCLC lines and engineered multiple pairs of isogenic lines with or without functional STK11/LKB1, we were able to test the relevance of GDF15 and its cellular (tumor versus host) sources in regulating cachexia development. These studies demonstrated that GDF15 secreted from *STK11/LKB1*-mutant tumors and not host tissue was critical for cachexia development. Using either neutralizing antibody or silencing of *GDF15* in tumor cells, tumor cell-derived GDF15 was relevant to wasting development and abrogation without affecting tumor progression, regardless of the systemic levels of GDF15. Interestingly, even in models with GDF15 serum concentrations at or below non-cachexia-inducing control models, we could still abrogate cachexia development with GDF15 neutralizing antibodies or silencing of tumor *GDF15* mRNA levels. There could be several explanations for this finding: 1) The ELISA we used could not detect small, but still clinically meaningful GDF15 levels, 2) At very low concentrations, GDF15’s effects were peripheral and not central, 3) Local increases in GDF15 could act on peripheral neurons to effectuate changes in wasting, and/or 4) Neutralization of GDF15 has effects on another molecule that was relevant to wasting in mice bearing STK11/LKB1-mutated tumors. In these model conditions, it will be critical to determine whether GDF15 is acting peripherally or centrally to promote wasting. We plan to use syngeneic model systems to characterize *STK11/LKB1*-mutant tumor-derived GDF15 central and peripheral effects leveraging tissue-specific knockout models of the GDF15 receptor, GFRAL. A more thorough evaluation of food intake with use of metabolic cages will also better demonstrate GDF15 sites of action. It is important to note that any peripheral effect on wasting by GDF15 could be translated from its actions directly on host tissues or indirectly through a central mechanism, i.e. the latter through GDF15’s central induction of hormones that then act peripherally.^26^

Furthermore, with our model systems, we were able to fill in knowledge gaps with regard to tumor growth kinetics as a function of cachexia induction and vice versa. When we utilized the GDF15 neutralizing antibody in our cachexia studies, we were uniformly able to suppress wasting without seeing a significant change in tumor growth kinetics, suggesting that growth of STK11/LKB1-mutated NSCLC cachexia-inducing tumors was not dependent on wasting by-products to profit through proliferation. Therefore, in our studies of cachexia, the growth of tumors with at least *STK11/LKB1* loss of function variants is not sensitive to wasting output. This cannot be extrapolated to tumors that grow secondary to other mutations or genetic drivers and is an area of active research. Finally, our studies also suggest a divergence in mechanisms downstream of *STK11/LKB1* loss of function with respect to regulation of tumor cell proliferation and cachexia wasting. Reconstitution of wild-type *STK11/LKB1* in H1437 tumor cells suppressed both tumor growth and cachexia induction. However, GDF15 neutralization or silencing in multiple lines only abrogated cachexia and not tumor growth suggesting a divergence of the STK11/LKB1-mutated cancer progression and cachexia pathways in tumor cells.

Any *in vivo* studies of GDF15 would be remiss without a thorough understanding of alterations in food intake with time, tumor development, and cachexia induction. We conducted longitudinal evaluations of food intake with all our studies. Interestingly, some cohorts showed a consistent and expected change in food intake, i.e. a decrease in food intake, associated with elevated GDF15 serum concentrations when comparing experimental groups vs controls. Our studies further illustrated that when the GDF15 increases in serum concentrations were not as high, there was less effect on food intake. We could explain these findings with one of several hypotheses. With smaller increases in GDF15 serum concentrations, there might not have been enough of a signal to centrally regulate food intake but still enough signal to promote wasting peripherally at the level of adipose and muscle. Alternatively, it could be our studies were not sensitive enough to pick up subtle changes in mouse food intake. To find subtle changes, these studies will need to be conducted in metabolic cages with mice singly-housed to optimize our detection of any food intake decrement with any increase in circulating GDF15 concentrations.

Finally, our previous studies employed human NSCLCs in immunodeficient mouse models. The lack of a complete immune system could have affected many findings – tumor growth, food intake, adipose wasting, muscle wasting, etc. To assess such immune system effects, we needed to study the same questions using isogenic murine tumor lines in syngeneic mice. In this manuscript, we obtained murine KP and KPL tumor lines previously derived from GEMMs^22^, permitting us to verify that the biologic link between *STK11/LKB1*-mutated tumors, GDF15 induction, and cachexia development was still intact in immunocompetent host backgrounds. To explore host versus tumor contributions in these syngeneic models, we will need to simultaneously conduct single cell studies to identify tumor-derived and host-derived changes. These models will provide information on not just GDF15, but other TGFβ family members and other cytokines associated with cachexia development.

Collectively, our work in this paper highlights the dependence of tumor cells with *STK11/LKB1* loss-of-function variants on increasing levels of GDF15 to promote cachexia, through either the increase of transcript, processing, and/or secretion of the protein. In our model systems, the tumor cells and not the host tissue are the source of GDF15 and its associated wasting. Both silencing of *GDF15* in tumor cells or use of a GDF15 neutralizing antibody systemically in multiple patient-derived and mouse-derived lines suppress STK11/LKB1-mutated NSCLC cachexia regardless of circulating GDF15 concentrations. Clinical translation of these findings will require testing of the ∼15% NSCLC patients whom possess *STK11/LKB1* variants to determine if the cachexia benefit from GDF15 inhibitors can similarly be observed across a wide range of systemic/circulating GDF15 levels.

## METHODS

### Materials

Detailed material information is listed in **Supplemental Table 1**.

### Buffer and Culture Medium

Buffer A contained 10 mM Tris-HCl (pH 6.8), 100 mM NaCl, 1% (w/v) SDS, 1 mM EDTA, 1 mM EGTA, Phosphatase Inhibitor Cocktail Set I and Set II, and Protease Inhibitor Cocktail. Medium A was RPMI 1640 supplemented with 5% (v/v) fetal bovine serum (FBS), 100 units/ml penicillin, and 100 µg/ml of streptomycin. Medium B was DMEM high glucose supplemented with 100 units/ml penicillin and 100 μg/ml of streptomycin. Medium C was medium B supplemented with 10% (v/v) FBS. Medium D was RPMI 1640 without glucose supplemented with 0.5% FBS, 100 units/ml penicillin, and 100 µg/ml of streptomycin.

### Cell Culture

Stock cultures of all human NSCLC cell lines (kind gift from John Minna), KP, KPL cell line (kind gift from Esra Akbay), and HEK293T cells (ATCC) were maintained in monolayer culture at 37°C in 5% CO_2_ in medium A or medium C, respectively. Each cell line was propagated, aliquoted, and stored under liquid nitrogen. Aliquots of these cell lines were passaged for less than 4 weeks to minimize genomic instability prior to injection into mice. Every six months, the cell lines were tested for *Mycoplasma* contamination using the MycoStrip *Mycoplasma* Detection Kit.

### Generation of Gene Knockout CRISPR/Cas9 Lentivirus

A single guide RNA (sgRNA) directed to *GFP* as non-targeting control (*ΔGFP*) or to human *GDF15* (*ΔGDF15*) were cloned into the *Bsm*BI (Thermo, #ER0451) site of lentiCRISPRv2-puro vector (AddGene, #52961) using the following pairs of annealed oligonucleotides obtained from IDT for GFP (forward 5’-CACCGGTGAACCGCATCGAGCTGA-3’; reverse 5’-AAACTCAGCTCGATGCGGTTCACC-3’), or human *GDF15* (forward 5’-CACCGGGACGTGACACGACCGCTG-3’; reverse 5’-AAAC CAGCGGTCGTGTCACGTCCC-3’). These constructs were transformed into stable competent *E. Coli* (NEB, #C3040H) and constructs were purified using the manufacture’s instruction of the MIDI-Prep system (Macherey-Nagel, #740410.10). The cloned nucleotides were confirmed by sanger sequencing. For lentiviral production, HEK293T cells were seeded in 100 mm dishes at ∼1 x 10^6^ cells in 10 ml medium D on day 0. On day 2 (∼70-80% confluence), medium was then removed and replaced with 8 ml medium C supplemented with an additional 2 mM glutamine, and 15 μl X-tremeGENE HP DNA Transfection Reagent, 2.5 μg of the indicated sgRNA-lentiCRISPRv2, 1 μg of pMD2.G (AddGene, #12259), and 1.5 μg of psPAX2 (AddGene, #12260) mixed in 0.8 ml Opti-MEM medium (Thermo, # 31985070). On day 3, medium was removed and treated with 11 ml medium B with 30% FBS supplemented with an additional 2 mM glutamine. On day 4, 5, and 6, virus-containing medium was collected and exchanged with 11 ml of fresh medium B with 30% FBS supplemented with an additional 2 mM glutamine. The collected virus-containing medium was filtered through a 0.45 μm syringe filter unit (Millipore Sigma) and stored at −80°C for future use.

### Generation of Wild-type STK11/LKB1 Overexpression Lentivirus

Complimentary DNA library was derived from the reverse transcription (Thermo, #N8080234) of extracted RNA from the H1792 line as described by the manufacturer’s directions as previously described. *STK11* (NM_000455.5) complimentary DNA containing the EcoRI and BamHI restriction sites at the 5’ and 3’ ends, respectively, using the following pairs of primers (forward 5’-TATTTCCGGTGAATTCatggaggtggtggacc-3’; reverse 5’-GAGAGGGGCGGGATCCtcactgctgcttgcag-3’). This fragment was cloned into the pLVX-EF1α-IRES-Puro vector (Clontech, #631988) by first treating with the EcoR1-HF (NEB, #R3101S) and BamH1-HF (NEB, #R3136S) restriction enzymes followed by In-Fusion Snap Assembly (Takara, #638947) using the manufactures’ directions. These constructs were transformed into stable competent *E. Coli* (NEB, #C3040H) and constructs were purified using the manufacture’s instruction of the MIDI-Prep system (Macherey-Nagel, #740410.10) The cloned nucleotides were confirmed by sanger sequencing. For lentiviral production, HEK293T cells were seeded in 100 mm dishes at ∼1 x 10^6^ cells in 10 ml medium D on day 0. On day 2 (∼70-80% confluence), medium was then removed and replaced with 8 ml medium C supplemented with 2 mM glutamine, and 15 μl X-tremeGENE HP DNA Transfection Reagent, 2.5 μg of the indicated pLVX-EF1α-IRES-Puro construct, 1 μg of pMD2.G (AddGene, #12259), and 1.5 μg of psPAX2 (AddGene, #12260) mixed in 0.8 ml Opti-MEM medium (Thermo, # 31985070). On day 3, medium was removed and treated with 11 ml medium B with 30% FBS supplemented with an additional 2 mM glutamine. On day 4, 5, and 6, virus-containing medium was collected and exchanged with 11 ml of fresh medium B with 30% FBS supplemented with an additional 2 mM glutamine. The collected virus-containing medium was filtered through a 0.45 μm syringe filter unit (Millipore Sigma) and stored at −80°C for future use.

### Generation of Knockout or Wild-type STK11/LKB1 Overexpression in Patient-derived NSCLC Lines

Monoclonal NSCLC cell lines were seeded in 6-well dishes at ∼1 x 10^5^ cell density in 2 ml of medium A. After 24 hours the medium was replaced with fresh medium A (non-transduced control) or 1 ml medium A and 1 ml of the indicated virus-containing medium, supplemented with 8 µg/ml polybrene reagent (Millipore Sigma, #TR-1003). After 24 hours, the medium was removed and replaced with 2 ml of medium A. The next day, the medium was removed and replaced with fresh 2 ml medium A containing 1 µg/ml puromycin, and puromycin-supplemented medium A was refreshed every 2 days until complete cell death was observed in the non-transduced control. Limiting dilution of these puromycin-selected cells was performed in puromycin-containing medium to produce single colonies of transduced cells, and these monoclonal knockout lines were subsequently propagated and aliquoted for storage under liquid nitrogen or further propagated for experiments described herein.

### Cancer Cachexia Animal Models

All animal studies were conducted under an Institutional Animal Care and Use Committee (IACUC)-approved protocol at UT Southwestern Medical Center (Dallas, Texas). NOD.CB17-Prkdc<scid>/J (*NOD/SCID*) and C57BL/6J mice were obtained from The Jackson Laboratory at approximately 9 weeks of age. All mice were allowed to acclimate in UT Southwestern animal facilities for at least 2 weeks. Animals were kept in a temperature-controlled facility (at approximately 22°C) with a 12 h light/dark cycle and were fed regular chow diet.

On day 0 of the study, *NOD/SCID* or C57BL/6J mice underwent baseline assessments of body weight (digital Ohaus scale) and both lean and fat mass components of body composition using ECHO MRI (ECHO Medical Systems). Then, 200 µl PBS in the absence or presence of either 1 x 10^7^ human or 1 x 10^6^ mouse NSCLC cells (parental or genetically engineered lines) were injected into the right flank of the *NOD/SCID* or C57BL/6J mice, respectively. Food intake measurements were conducted daily as described previously, including the method for calculation of average daily food intake.^27^ Every 2-3 days, mice underwent longitudinal measurements of body weight, both lean and fat mass components of body composition, and tumor size by caliper (VWR) measurements of length, width, and breadth.

The timing of animal sacrifice for tumor-injected cohorts was a function of imminent animal death (expected within 12 hours), tumor volumes approaching 2.0 cm diameter, or cohorts of animal reaching about 50% fat loss; PBS cohorts were sacrificed concurrently with matched tumor cohorts. At sacrifice, mice were euthanized as recommended by the IACUC with use of a CO_2_ chamber. Tumor weights were measured and collected on dry ice prior to long-term storage at −80°C (for future analysis). Whole blood at sacrifice was obtained through cardiac puncture and serum was obtained by subjecting the whole blood to centrifugation at 960 x g at 4°C for 10 min followed by removal of the supernatant.

Longitudinal values for tumor volume were calculated as half the product of caliper measurements of length, width, and breadth. Longitudinal and terminal values for body weight, lean mass, and fat mass were calculated relative to both the matched animal day zero value and mean value of the matched PBS cohort. At sacrifice, relative tumor-free body weight and lean mass were calculated by subtracting the matched animal tumor weight at sacrifice prior to normalization.

### Serum GDF15 Protein Concentration

Human GDF15 in plasma was measured using a human GDF15 Quantikine ELISA kit (R&D Systems, #DGD150,) following the manufacturer’s instructions. Mouse GDF15 in plasma was measured using a mouse/rat GDF15 Quantikine ELISA kit (R&D Systems, #MGD150) following the manufacturer’s instructions.

### Immunoblot Analysis

Human NSCLC cell lines cells were seeded in 6-well dishes at ∼4 x 10^5^ cell density in 2 ml of medium A. After 24 hours cells were washed with 2 ml DPBS, then treated with Medium A or Medium D in the absence or presence of 10 mM glucose. After 16 hours, media were harvested and spun with 800 *g* for 2 min and the supernatant was concentrated 15-fold using a 3 kDa filter (Amicon Ultra) per manufacturer’s instructions to obtain the concentrate. Cells were washed with 2 ml DPBS, then cells were treated with 170 μl of buffer A then subjected to a plate shaker for 2 min at 700 RPM followed by harvesting of the cell lysate which was subjected to 10 s of sonication (Cole-Parmer, #EW-04714-50) at 50% amplitude. The concentrated media and cell lysate were then used for immunoblot analysis.

Tumor from mouse studies were thawed on ice and approximately 50-60 mg of tumor tissue was placed in a 2 ml microtubes with 1 ml of Buffer A and homogenized with a Bead Ruptor Elite apparatus (Omni International) for three cycles of 20 s at 4 m/s. The homogenized sample was immediately subjected to centrifugation at 16°C for 10 min at 20,000 x g. The supernatant was separated from the pellet and collected, followed by 10 s of sonication (Cole-Parmer, #EW-04714-50) at 50% amplitude. The tissue lysate or mouse serum were then used for immunoblot analysis.

Protein concentrations of cell medium, cell lysate, mouse serum, or mouse tumor lysate were measured using a Pierce^TM^ bicinchoninic acid kit and then mixed with 5X loading buffer (250 mM Tris-HCl (pH 6.8), 10% sodium dodecyl sulfate, 25% glycerol, 5% β-mercaptoethanol, and 0.2% bromophenol blue) heated at 95°C for 5 minutes, and subjected to 10% or 15% SDS/PAGE (8 µg/lane for cell/tumor lysate, equal volume for serum, or volume of cell medium normalized to cell protein concentration). The electrophoresed proteins were transferred to nitrocellulose membranes using the Bio-Rad Trans Blot Turbo system, followed by incubation in blocking buffer containing 5% (w/v) non-fat powdered milk in PBS-T. Blocked membranes were washed briefly in PBS-T, followed by primary staining with indicated primary antibodies diluted in primary antibody solution-Can Get Signal (TOYOBO). Membranes were again washed in PBS-T, followed by secondary staining with donkey anti-mouse IgG conjugated to horseradish peroxidase diluted in the blocking buffer. Membranes were again washed in PBS-T, and bound antibodies were visualized by brief incubation in chemiluminescent substrate solution followed by imaging using an Odyssey FC Imager Dual-mode Imaging System (2-min integration time). Immunoblot images were analyzed using Image Studio software (LICOR).

#### Real-time PCR Analysis of Gene Expression

Extraction of total RNA with RNA-STAT-60 reagent from tumors and subsequent quantitative reverse transcriptase PCR assays for mRNA levels of the indicated gene products were conducted as previously described (*1*). The sequences of primers (obtained from Integrated DNA Technologies) used in these studies are as follows:

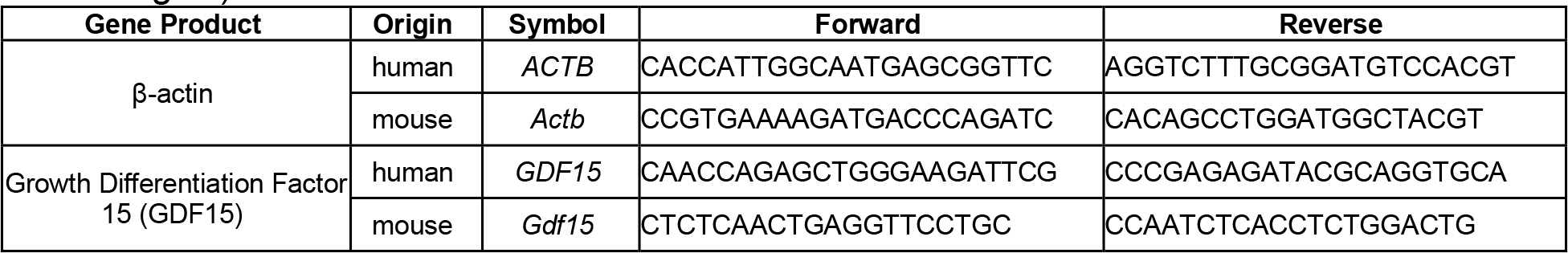

#### Microscopic Analysis of Mouse Adipose and Muscle Tissue

Epididymal white fat was collected, and immersion fixed in 10% neutral-buffered formalin. All samples were paraffin processed by members of UT Southwestern’s Histo Pathology Core using standard histologic technique.^28, 29^ Resulting embeds were sectioned at 5 µm, and produced slides were stained by regressive methodology utilizing Leica Selectech reagents (Hematoxylin 560, Define, Blue Buffer, and Eosin-Y 515) on a Tissue-Tek Prisma Plus robotic stainer (Sakura-USA). From the stained slides, microscopy fields were imaged on a BZ-X scanner (Keyence) at 20X magnification and adipocyte area was manually measured using ImageJ software (NIH) for three fields from each slide (250-350 adipocytes per slide). Each group contains the quantification from three slides prepared from three independent mice. Publication proofs of representative fields were obtained on the Keyence BZ-X scanner at 20X magnification.

Whole gastrocnemius skeletal muscle samples were obtained from the limb contralateral to the tumor-implanted flank and cryo-embedded in transverse myofiber orientation protruding from a 1:8 volume mixture of Gum Tragacanth powder (Sigma-Aldrich) to Tissue Freezing Medium (TFM; Triangle Bioscience). All embeds were snap frozen in an isopentane heat extractant bath supercooled to freezing temperature of −155°C in liquid nitrogen. Resulting blocks were stored at −80°C prior to sectioning. Subsequently, gastrocnemius cryoblocks were equilibrated to cutting temperature in a Leica CM3050S cryostat and eight-micron sections were prepared from the transverse face of each block (−25°C object temperature, −23°C chamber temperature, 3.5° blade angle). Slides were briefly air-dried and returned to −80°C for metachromatic ATPase staining at a later point in time.

Gastrocnemius was collected and calcium mediated metachromatic ATPase staining was performed according to previously reported methods.^30–32^ In detail, resulting slides were incubated at room temperature with agitation for eight minutes in pH 4.4 ATPase-inhibiting differentiation buffer, containing 0.05M Potassium Acetate and 0.018M Calcium Chloride Dihydrate. Slides were rinsed free of differentiation buffer and returned to neutral pH by brief incubation in three sequential pH 7.8 acid-neutralizing baths containing 0.1M Tris and 0.018M Calcium Chloride Dihydrate. Next, slides were incubated in ATPase reactant buffer for 25 minutes at room temperature with agitation. pH 9.4 ATPase reactant buffer, comprised of 0.05M Glycine, 0.03M Calcium Chloride Dihydrate, 0.06M Sodium Chloride, was prepared immediately before use. Following ATPase reactant buffer incubation, slides were briefly rinsed in three changes of 1% Calcium Chloride Dihydrate before staining with aqueous 0.1% Toluidine Blue for one minute. Following colorization with Tol-Blue, slides were briefly rinsed in two changes of distilled water, differentiated in 95% ethanol, dehydrated through three changes of 100% ethanol and cleared in xylene. All mentioned chemical salts were sourced from Millipore-Sigma (St. Louis, MO). Ethanol and xylene were sourced from Greenfield Global (Shelbyville, KY).

Coverslips were applied with Permount synthetic mounting media (Fisher Scientific). From the stained slides, microscopy fields from the anterior aspect of each gastrocnemius’ medial head were imaged using a Keyence BZ-X microscope at 10X magnification for morphometric analysis. Type I and Type II myofibers were distinguished based on the metachromatic ATPase staining. Myofibers were outlined from each anterior-medial gastrocnemius and their myofiber cross-sectional area manually measured in ImageJ. Each group contains the quantification of a total of 150-300 myofibers collected from three independent mice.

#### Statistical Analysis

Details of statistical analysis for each experiment can be found in the respective figure legend. Data is presented as mean ± SEM, dot plots ± SEM, dot plots with bars ± SEM, or histogram. For experimental designs with two conditions, unpaired parametric *t*-tests were conducted to investigate significant differences following tests to verify normality and equal variance of both groups. For experimental designs with greater than two conditions, one-or two-way analysis of variance (ANOVA) tests were conducted to examine if there were significant differences in outcomes among groups. For the case of multiple pairwise comparisons, Dunnett’s multiple comparison post-test was used. All statistical analyses were computed using GraphPad Prism 10 software.

### Study Approval

Animal studies were approved through the University of Texas Southwestern Medical Center’s Institutional Animal Care and Use Committee (protocol 2015-100994).

## Abbreviations

(CC): cancer cachexia
(CRC): colorectal cancer
(IO): immuno-oncology
GDF15: growth differentiation factor 15 (gene: *GDF15*)
(NSCLC): protein, non-small cell lung cancer
STK11 or LKB1: serine/threonine kinase 11/liver kinase B1 (gene: *STK11* or *LKB1*, protein:)

## ACKNOWLEDGEMENTS

We thank Michael Brown, Joseph Goldstein, Shawn Burgess, and Jay Horton for their suggestions and review. We also thank Bei Zhang (Pfizer), Mingjian Lu (Pfizer), Junjie Li (Pfizer), Ja Young Kim-Mueller (Pfizer) and Danna Breen (Pfizer) for their contributions. We thank Natalie Lopez and Min Ding for their excellent technical assistance. This work was supported by grants from the Burroughs Wellcome Fund Career Awards for Medical Scientists (1019692); American Cancer Society Grant (133889-RSG-19-195-01-TBE); Cancer Prevention and Research Institute of Texas (RP230140 and RP210041); and National Institutes of Health grants (1R01CA266900, 1P30DK127984, P30CA142543, P50CA070907, and T32DK007745).

---

**Supplemental Figure 1.**
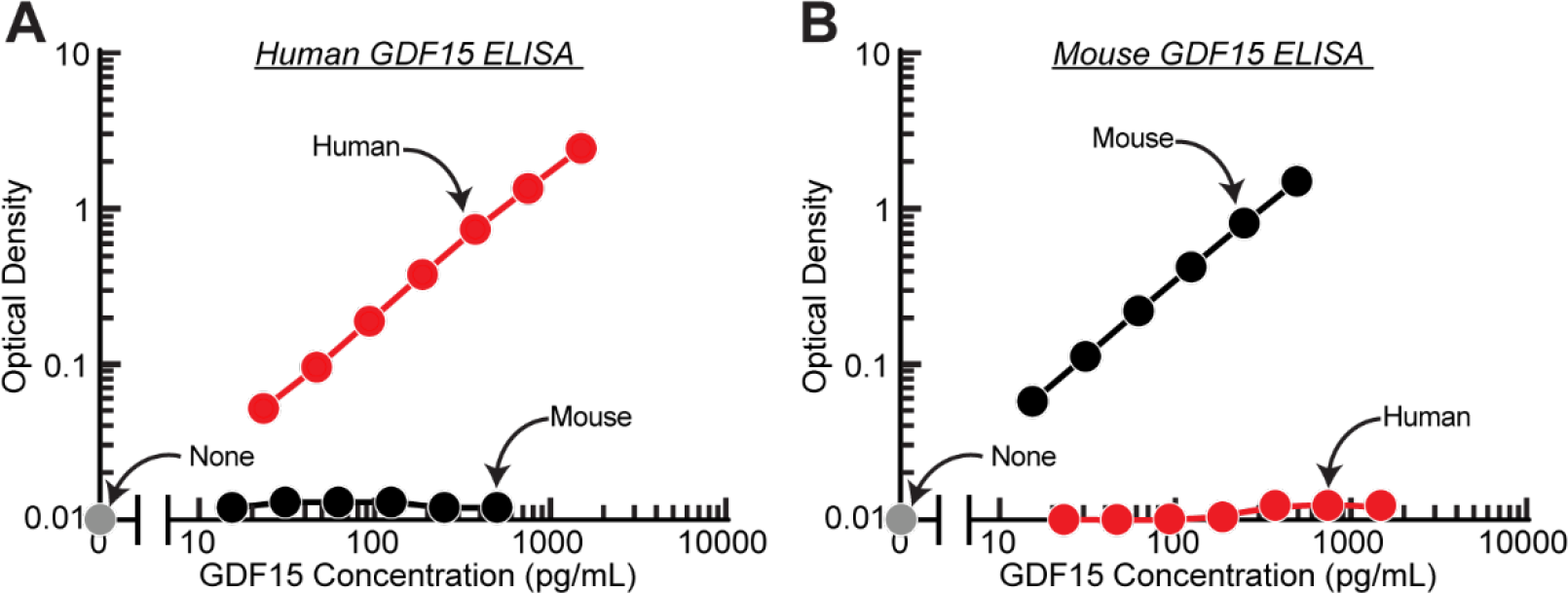
Specificity of Human and Mouse GDF15 ELISAs. A-B) The indicated concentration of human (red) or mouse (black) GDF15 standards were subjected to human (A) or mouse (B) ELISA per the manufacturer’s instructions.

**Supplemental Figure 2.**
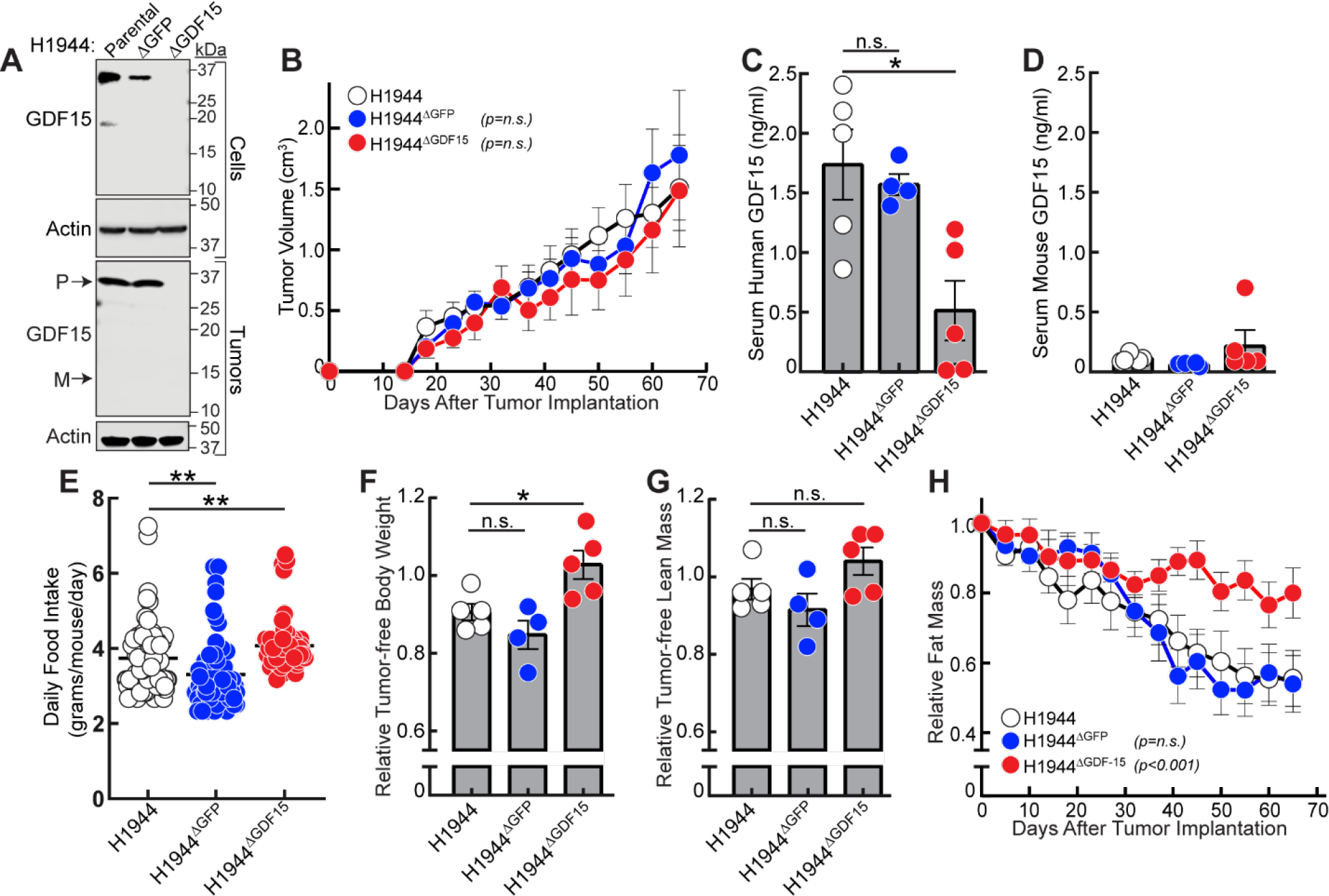
G*D*F15 silencing in *STK11/LKB1*-mutated NSCLC tumors allotransplanted into mice that have intermediate circulating GDF15 concentrations. A-K) Chow-fed 15-week-old NOD/SCID male mice (n = 5-6 per group) were injected s.c. with 200 ul PBS in the absence or presence of 1 x 10^7^ cells from parental H1944, H1944^ΔGFP^, or H1944^ΔGDF15^ lines as described in Methods. Longitudinal or endpoint measurements of tumor volume (B), food intake (E), tumor-free body weight (F), tumor-free lean mass (G), or fat mass (H) were obtained as described in Methods. Tumor cells before injection and tumors at sacrifice were processed for immunoblot analysis with the indicated antibody (A). Serum at sacrifice was subjected to the human (C) or mouse (D) GDF15 ELISA as described in Methods. Data are shown as mean ± SEM of the actual measurements (B-E) or relative to their day 0 values (F-H). P was calculated using 1-way (C-G) or 2-way (B, H) ANOVA followed by Dunnett’s multiple-comparison test for significant differences from the H1944 parental cohort. *\*p<0.05; **p<0.01; n.s. = not significant; M = GDF15 mature protein; P = GDF15 proprotein*.

**Supplement Figure 3.**
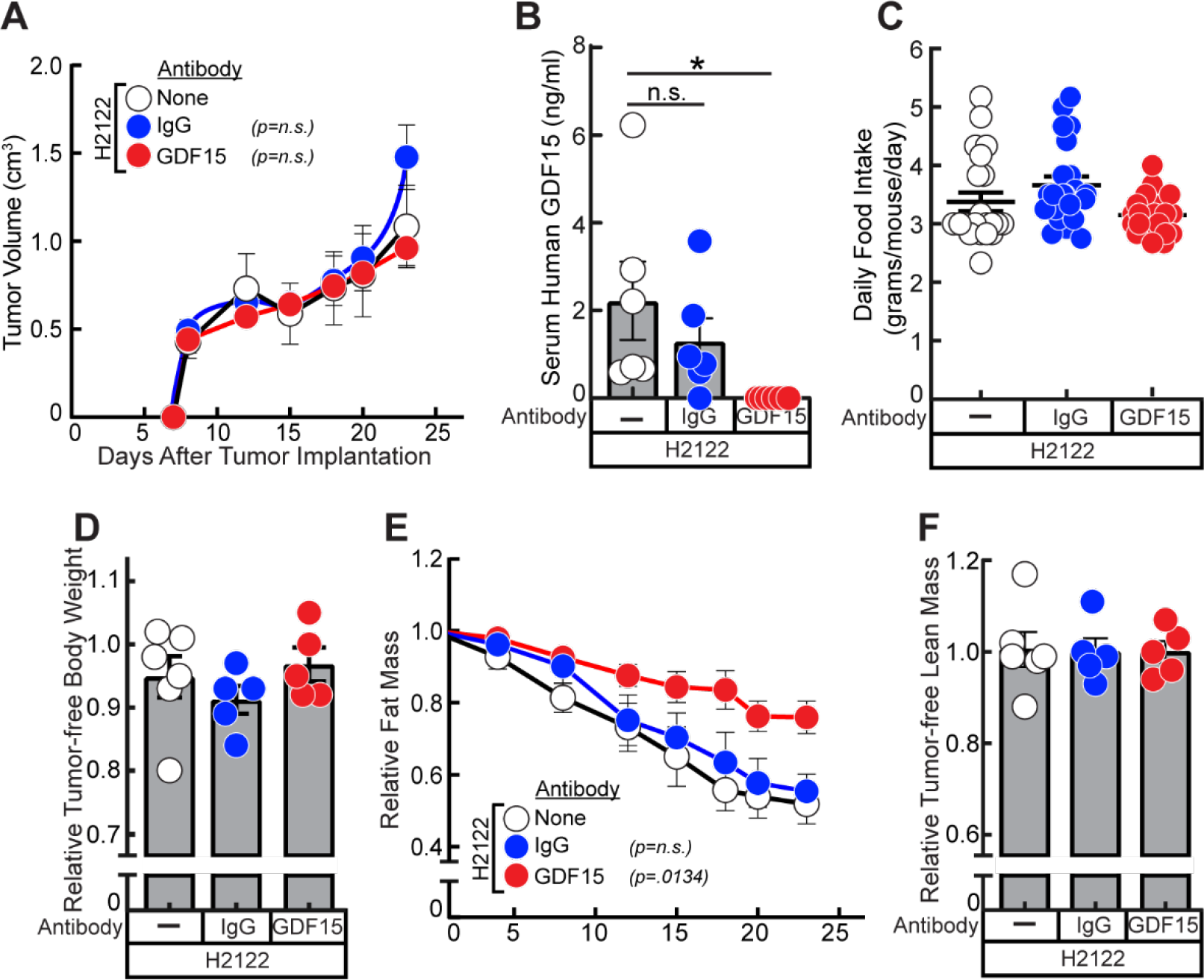
GDF15 antibody neutralization in mice bearing *STK11/LKB1*-mutated NSCLC tumors with low circulating GDF15 concentrations. A-F) Chow-fed 14-week-old NOD/SCID male mice (n = 6) were injected s.c. with 200 μl PBS containing 1 x 10^7^ cells from parental H2122 human NSCLC line on day 0. Starting on day7, mice were injected with s.c. 150 μl PBS in the absence or presence of 10 mg/kg IgG or GDF15 antibody weekly. Longitudinal or endpoint measurements of tumor volume (A), food intake (C), tumor-free body weight (D), fat mass (E), and tumor-free lean mass (F) were obtained as described in Methods. Serum at sacrifice was processed for human GDF15 ELISA (B) as described in Methods. Data are shown as mean ± SEM of the actual measurements (A-C) or relative to their day 0 values (D-F). P was calculated using 1-way (B, C, D, F) or 2-way (A, E) ANOVA followed by Dunnett’s multiple-comparison test for significant differences from the H2122 parental cohort. *p**** <0.0001; n.s. = not significant; M = GDF15 mature protein; P = GDF15 proprotein*.

**Supplemental Figure 4.**
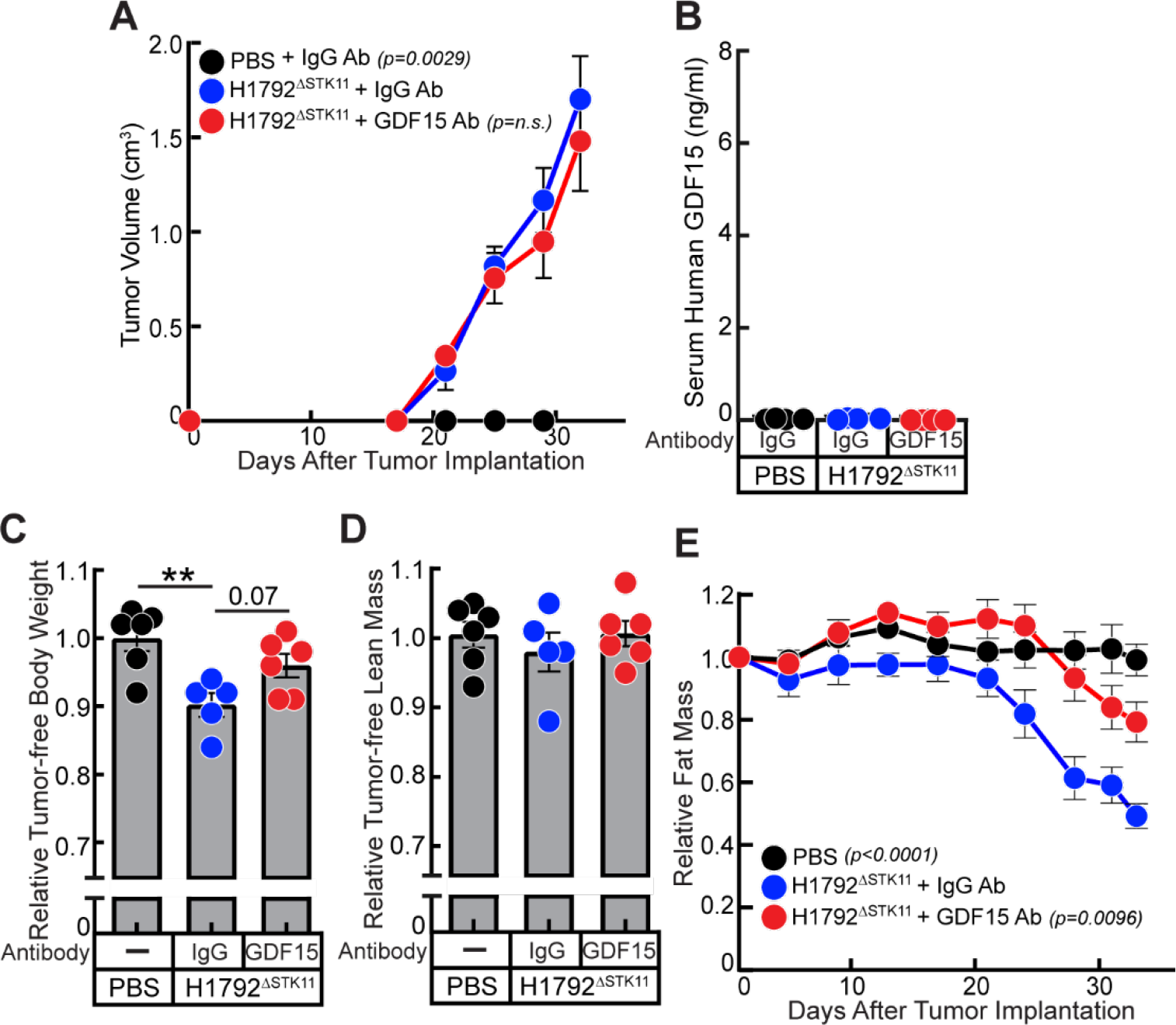
Dependence of mice bearing *STK11/LKB1*-silenced NSCLC isogenic tumors on GDF15 for cachexia induction. A-E) Antibody neutralization of *STK11/LKB1*-silenced NSCLC model. Chow-fed 12-week-old NOD/SCID male mice (n = 6) were injected s.c. with 200 μl PBS containing in the absence or presence of 1 x 10^7^ cells from H1792^ΔSTK11^ human NSCLC line on day 0. Starting on day 7, mice were injected with s.c. 150 μl PBS in the absence or presence of 10 mg/kg IgG or GDF15 antibody weekly. Longitudinal or endpoint measurements of tumor volume (A), tumor-free body weight (C), tumor-free lean mass (D), and fat mass (E) were obtained as described in Methods. Serum was processed for immunoblot analysis for human (B) GDF15 ELISA as described in Methods. Data are shown as mean ± SEM of the actual measurements (A-B) or relative to their day 0 values (C-E). P was calculated using 1-way (B-D) or 2-way (A, E) ANOVA followed by Dunnett’s multiple-comparison test for significant differences from the H1792^ΔSTK11^ treated with IgG antibody cohort. *p** <0.01; n.s. = not significant*.

**Supplemental Figure 5.**
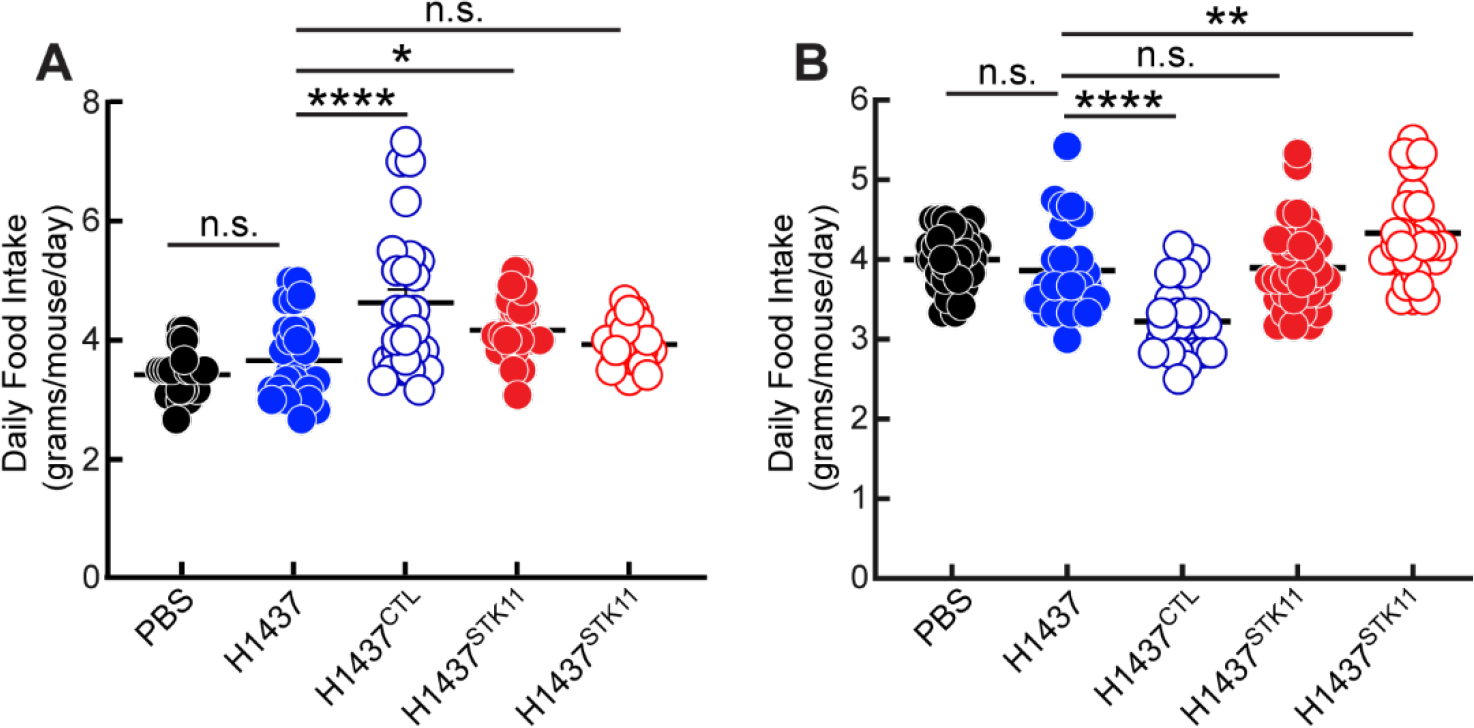
Effect of food intake on wild-type STK11/LKB1-reconstituted tumor-bearing mice. A-B) Chow-fed 17-week-old (A) or 15-week-old (B) NOD/SCID male mice (n = 6 per group) were injected s.c. with 200 ul PBS in the absence or presence of 1 x 10^7^ cells from parental H1437, H1437^CTL^, H1437^STK11^, or another H1437^STK11^ monoclonal line as described in Methods. Longitudinal measurements of food were obtained over two separate experiments as described in Methods. Data are shown as mean ± SEM of the actual measurements. P was calculated using 1-way ANOVA followed by Dunnett’s multiple-comparison test for significant differences from the parental H1437 cohort. *\*p<0.05; **p<0.01; p****<0.0001; n.s. = not significant*.

## SUPPLEMENTAL TABLES

**Supplemental Table 1.**
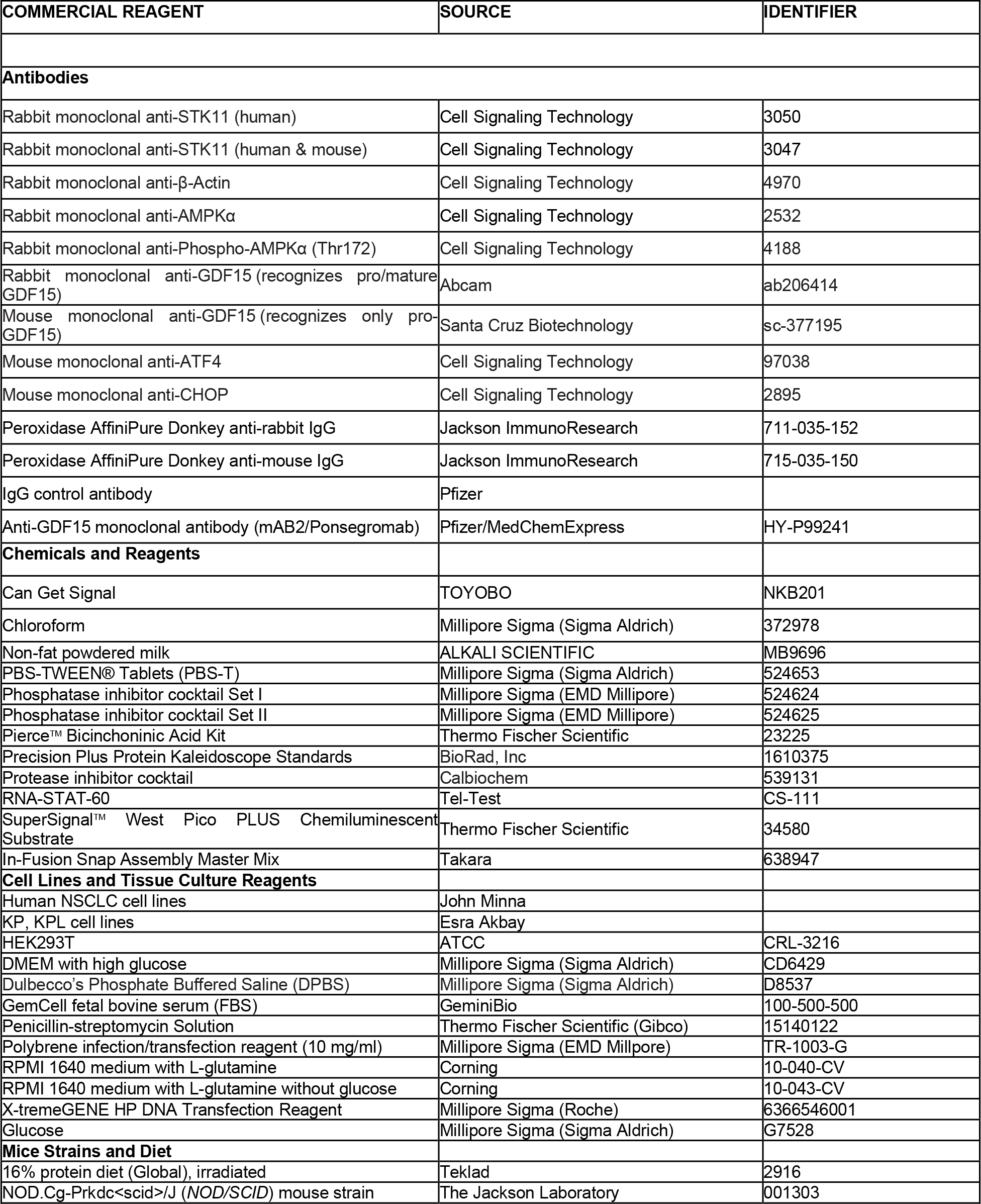
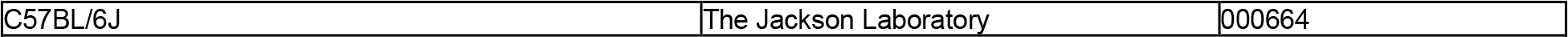
Materials for Assays.

| COMMERCIAL REAGENT | SOURCE | IDENTIFIER |
| --- | --- | --- |
| **Antibodies** |  |  |
| Rabbit monoclonal anti-STK11 (human) | Cell Signaling Technology | 3050 |
| Rabbit monoclonal anti-STK11 (human & mouse) | Cell Signaling Technology | 3047 |
| Rabbit monoclonal anti-β-Actin | Cell Signaling Technology | 4970 |
| Rabbit monoclonal anti-AMPKα | Cell Signaling Technology | 2532 |
| Rabbit monoclonal anti-Phospho-AMPKα (Thr172) | Cell Signaling Technology | 4188 |
| Rabbit monoclonal anti-GDF15 (recognizes pro/mature GDF15) | Abcam | ab206414 |
| Mouse monoclonal anti-GDF15 (recognizes only pro-GDF15) | Santa Cruz Biotechnology | sc-377195 |
| Mouse monoclonal anti-ATF4 | Cell Signaling Technology | 97038 |
| Mouse monoclonal anti-CHOP | Cell Signaling Technology | 2895 |
| Peroxidase AffiniPure Donkey anti-rabbit IgG | Jackson ImmunoResearch | 711-035-152 |
| Peroxidase AffiniPure Donkey anti-mouse IgG | Jackson ImmunoResearch | 715-035-150 |
| IgG control antibody | Pfizer |  |
| Anti-GDF15 monoclonal antibody (mAB2/Ponsegromab) | Pfizer/MedChemExpress | HY-P99241 |
| **Chemicals and Reagents** |  |  |
| Can Get Signal | TOYOBO | NKB201 |
| Chloroform | Millipore Sigma (Sigma Aldrich) | 372978 |
| Non-fat powdered milk | ALKALI SCIENTIFIC | MB9696 |
| PBS-TWEEN® Tablets (PBS-T) | Millipore Sigma (Sigma Aldrich) | 524653 |
| Phosphatase inhibitor cocktail Set I | Millipore Sigma (EMD Millipore) | 524624 |
| Phosphatase inhibitor cocktail Set II | Millipore Sigma (EMD Millipore) | 524625 |
| Pierce™ Bicinchoninic Acid Kit | Thermo Fischer Scientific | 23225 |
| Precision Plus Protein Kaleidoscope Standards | BioRad, Inc | 1610375 |
| Protease inhibitor cocktail | Calbiochem | 539131 |
| RNA-STAT-60 | Tel-Test | CS-111 |
| SuperSignal™ West Pico PLUS Chemiluminescent Substrate | Thermo Fischer Scientific | 34580 |
| In-Fusion Snap Assembly Master Mix | Takara | 638947 |
| **Cell Lines and Tissue Culture Reagents** |  |  |
| Human NSCLC cell lines | John Minna |  |
| KP, KPL cell lines | Esra Akbay |  |
| HEK293T | ATCC | CRL-3216 |
| DMEM with high glucose | Millipore Sigma (Sigma Aldrich) | CD6429 |
| Dulbecco's Phosphate Buffered Saline (DPBS) | Millipore Sigma (Sigma Aldrich) | D8537 |
| GemCell fetal bovine serum (FBS) | GeminiBio | 100-500-500 |
| Penicillin-streptomycin Solution | Thermo Fischer Scientific (Gibco) | 15140122 |
| Polybrene infection/transfection reagent (10 mg/ml) | Millipore Sigma (EMD Millipore) | TR-1003-G |
| RPMI 1640 medium with L-glutamine | Corning | 10-040-CV |
| RPMI 1640 medium with L-glutamine without glucose | Corning | 10-043-CV |
| X-tremeGENE HP DNA Transfection Reagent | Millipore Sigma (Roche) | 6366546001 |
| Glucose | Millipore Sigma (Sigma Aldrich) | G7528 |
| **Mice Strains and Diet** |  |  |
| 16% protein diet (Global), irradiated | Teklad | 2916 |
| NOD.Cg-Prkdc<sup>scid</sup>/J (NOD/SCID) mouse strain | The Jackson Laboratory | 001303 |
| C57BL/6J | The Jackson Laboratory | 000664 |

